# MgATP/MgADP-dependent conformational dynamics and intrinsically disordered regions of vascular K_ATP_ channels revealed by cryoEM

**DOI:** 10.64898/2026.09.28.753909

**Authors:** Camden M. Driggers, Yi-Ying Kuo, Zhongying Yang, Sun-Joo Lee, Colin G. Nichols, Katarzyna Walczewska-Szewc, Show-Ling Shyng

**Author notes:** To whom correspondence may be addressed: Camden M. Driggers,; Katarzyna Walczewska-Szewc,; Show-Ling Shyng.

## Abstract

Vascular smooth muscle K_ATP_ channels, composed of the pore-forming Kir6.1 and regulatory SUR2B subunits, control vascular tone, dysfunction of which causes systemic disease. Vascular K_ATP_ is regulated by Mg-nucleotides, but the underlying structural mechanism has remained elusive. Here, we determined cryoEM structures of these channels in the presence of MgATP and MgADP. Two key structures captured, one showing the SUR2B-nucleotide binding domains (NBDs) separated and one showing the SUR2B-NBDs dimerized, reveal conformation-specific organization of intrinsically disordered regions (IDRs) found in both Kir6.1 and SUR2B. In the NBD-separated conformation, the Kir6.1-N terminal IDR (KNt) sits within the central cleft of SUR2B’s ABC-core. In the NBD-dimerized conformation, KNt is excluded from the central cleft and instead forms contacts with an ED domain comprising 15 consecutive glutamate and aspartate residues within a SUR2B IDR, the N1-T2 linker connecting NBD1 (N1) to transmembrane domain 2 (T2). Moreover, within the N1-T2 linker a regulatory helix seen between the two NBDs in the NBD-separated conformation moves to outside the dimerized NBDs, interacting with the C-terminal residues unique to SUR2B, in the NBD-dimerized conformation. MD simulations further reveal that transient but frequent interactions mediated by the IDRs may facilitate Mg-nucleotide dependent conformational switch in vascular K_ATP_ channels.

## INTRODUCTION

ATP-sensitive potassium (K_ATP_) channels couple cell energetics with membrane excitability to govern physiological processes in response to metabolic changes^1^. In vascular smooth muscle, K_ATP_ channels formed by Kir6.1 and SUR2B control vascular tone^2–5^. Activation of vascular K_ATP_ leads to membrane hyperpolarization and vasodilation^6^. while inhibition causes membrane depolarization and vasoconstriction^5–8^. Gain-of-function mutations of vascular K_ATP_ channels cause Cantú syndrome^9,10^, a severe systemic hypotension disorder^2,11^.

In addition to vascular K_ATP_ (vK_ATP_), pancreatic K_ATP_ (pK_ATP_) comprising Kir6.2 and SUR1 and cardiac K_ATP_ comprising Kir6.2 and SUR2A (cK_ATP_) are two other major K_ATP_ channel isoforms^3,12^, which regulates insulin secretion in pancreatic endocrine β-cells and protects ventricular cardiac myocytes against ischemic injuries, respectively. Kir6.1 and Kir6.2 are members of the inwardly rectifying potassium (Kir) channel family, with two transmembrane helices M1 and M2 and cytoplasmic N- and C-terminal domains. SUR proteins, including SUR1, SUR2A and SUR2B, the latter two differing in their C-terminal 42 amino acids (C42) due to alternative splicing of the SUR2 gene, are members of the ABC protein family. Each SUR contains an N-terminal transmembrane domain (TMD0) and an ABC core of two transmembrane domains TMD1 and TMD2 and two cytoplasmic nucleotide binding domains (NBDs) NBD1 and NBD2 (Fig.1). While all three K_ATP_ isoforms are regulated by intracellular ATP and ADP, they have distinct gating properties and nucleotide sensitivities. Notably, unlike pK_ATP_ and cK_ATP_ which contain Kir6.2, vK_ATP_ containing Kir6.1 lacks spontaneous activity in the absence of inhibitory ATP and requires Mg-nucleotide acting on SUR2B for opening^2–4^.

**Figure 1.**
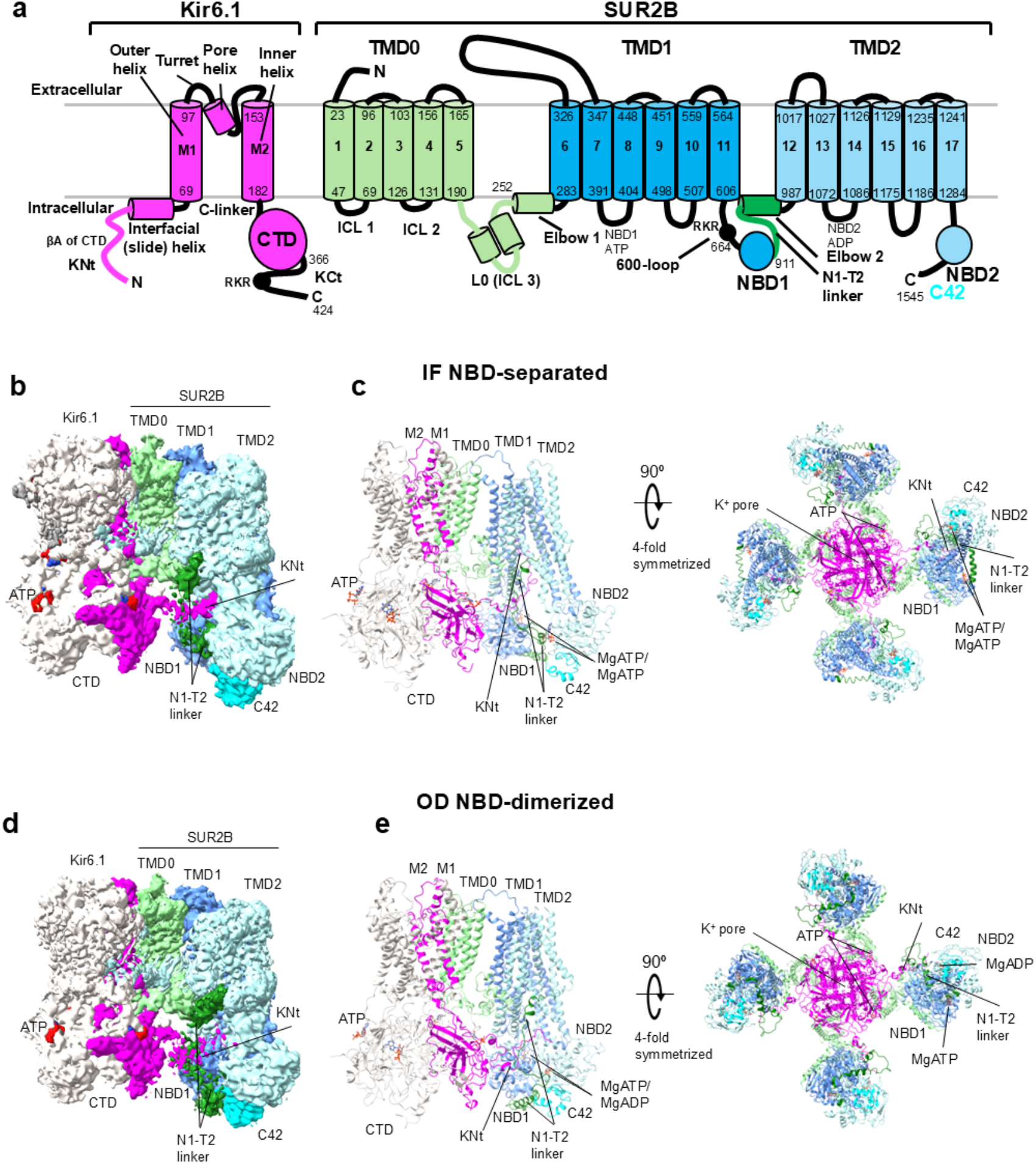
Discrete conformations of NBD-separated and NBD-dimerized structures of the Kir6.1/SUR2B K_ATP_ channels prepared in the presence of MgADP and MgATP. **(a)** Topology diagram of SUR2B and Kir6.1 domain organization including numbering of the inner and cytoplasmic loops, transmembrane helices, SUR2B cytoplasmic nucleotide binding domains (NBD1 and NBD2), the cytoplasmic domain of Kir6.1 (CTD), and intrinsically disordered regions (IDRs). IDRs are the N-terminal of Kir6.1 (KNt), the C-terminal of Kir6.1 (KCt), the TMD1-NBD1 loop (600-loop), and the NBD1 to TMD2 (N1-T2) linker. **(b)** CryoEM map of Kir6.1 tetrameric core and one SUR2B subunit in an inward-facing (IF) NBD-separated conformation shown in a side view, with the map for ATP colored red and other map features colored similarly to the topology diagram in (a), with three of the four Kir6.1 subunits colored light grey. **(c)** Structural model of Kir6.1 tetrameric core and one SUR2B subunit in an IF NBD-separated conformation shown from a side view (*left*) and a cytoplasmic view of a four-fold symmetrized structural model showing the full vascular K_ATP_ channel (*right*). **(d)** CryoEM map of the Kir6.1 tetrameric core and one SUR2B subunit in an NBD-dimerized occluded (OD) conformation. **(e)** Structural model of the Kir6.1 tetrameric core and one SUR2B subunit in an NBD-dimerized occluded (OD) conformation shown from a side view (*left*), and a cytoplasmic view of the full vascular K_ATP_ channel (*right*) shown like in (c).

High resolution cryoEM structures of K_ATP_ channels, mostly pK_ATP_, have shed light on the general molecular mechanisms of channel interaction and regulation by physiological and pharmacological ligands^13^. ATP and ADP control K_ATP_ activity via allosteric sites on both Kir6 and SUR. Mg^2+^-independent binding of ATP, and to a lesser extent ADP, to inhibitory sites in Kir6 closes the channel^1^. In contrast, Mg^2+^-complexed ATP and ADP (MgATP/MgADP) stimulate channel activity by induced dimerization of SUR’s two NBDs upon MgATP binding at NBD1 and MgADP binding at NBD2. Pharmacological inhibitors, exemplified by the sulfonylurea drug glibenclamide, bind in a transmembrane pocket above SUR-NBD1^14,15^, immobilize the N-terminal Kir6 (KNt) in the central cleft of SUR1’s ABC core adjacent the bound drug, thereby preventing MgATP/MgADP-induced NBD dimerization. In contrast, K_ATP_ openers bind at the interface of the two TM bundles of the SUR-ABC core, which is thought to stabilize MgATP/MgADP-induced NBD dimerization^16^. Despite the progress, our understanding of K_ATP_ structure-function relationships is far from complete. In particular, the structural bases underlying isoform-specific gating and nucleotide sensitivity are not clearly understood. Moreover, while NMR studies of isolated NBDs and surrounding linker sequence have implicated a modulatory role of the intrinsically disordered linker sequence in NBD function^17^, many intrinsically disordered regions (IDRs) in Kir6 and SUR are still unresolved in the context of the full K_ATP_ channel and their functional roles remain enigmatic.

Here, we report cryoEM structures of vK_ATP_ in the presence of MgATP and MgADP. From a single cryoEM sample we observed two distinct structures of the fully-assembled channel, including SUR2B in an NBD-separated conformation and one in an NBD-dimerized conformation. Significantly, these structures reveal the cryoEM densities of previously unseen IDRs of both Kir6.1 and SUR2B. All-atom molecular dynamics (MD) simulations of the full channel capture conformation-specific interactions between different IDRs and between IDRs and structured domains. Our structures together with MD simulations uncover a crucial role of IDRs in MgADP/MgATP-dependent conformational switch in vK_ATP_ channels with implications for isoform-specific gating regulation.

## RESULTS

### Structure determination of vK_ATP_ channels comprising Kir6.1 and SUR2B

Vascular K_ATP_ channels purified from COSm6 cells co-expressing rat Kir6.1 and SUR2B (97.6 and 97.2% sequence identity to human Kir6.1 and SUR2B, respectively) as independent polypeptides were used for cryoEM as described previously (see Methods)^14^. Two independent datasets were collected from purified vK_ATP_ incubated with 1 mM ATP, 1 mM ADP, 4 mM Mg^2+^, with or without 10 µM VU0542270, a recently reported vK_ATP_-selective inhibitor^18^ (Figs.S1, S2). Although VU0542270 was added to the first dataset sample, we did not observe non-protein cryoEM density that could be assigned as VU0542270, suggesting that the presence of MgATP/MgADP reduced VU0542270 binding, as has been shown for the well characterized inhibitor, glibenclamide^19^. Moreover, the cryoEM maps from both datasets are nearly identical, and both captured SUR2B in two major distinct conformations: NBD-separated or NBD-dimerized, indicating the structures obtained from both datasets reflect MgATP/MgADP-dependent regulation. Because the first dataset containing more particles generated maps with higher resolutions (Figs.S1, S2), it was used for subsequent structural analysis.

Two-dimensional (2D) class averages of 102,693 particles indicated an ordered symmetrical Kir6.1 and SUR2B-TMD0, with conformational heterogeneity of the SUR2B-ABC core especially the NBDs, which deviate from fourfold symmetry (Fig.S1). Initial cryoEM maps from CryoSPARC *Ab initio* reconstruction followed by C1 homogeneous refinement extended to 3.5 Å resolution by GSFSC (the Gold-Standard Fourier Shell Correlation) 0.143^20,21^. Heterogeneity of SUR2B limits the map quality for the NBDs. To circumvent this limitation, we applied fourfold symmetry expansion and focused three-dimensional (3D) classification^22^, using a mask that included the Kir6.1 tetramer core and a single SUR2B subunit (Kir6.1_4_SUR2B_1_; Fig.S1), yielding six conformational classes. Two classes showed the NBDs in separated conformation characteristic of type IV ABC transporters in the inward-facing (IF) conformation. Two showed NBDs dimerized resembling type IV ABC transporters in the occluded (OD) conformation. The other two classes had the NBDs in between the separated and dimerized classes, or NBD2 poorly resolved. Particles from the two IF classes were combined, and the OD class which refined to the highest resolution (3.5 Å; Fig.S1) underwent further local refinement focusing on the SUR2B subunit. This resulted in improved map quality for the NBDs albeit with lower nominal overall resolutions (Fig.S1). The NBD-focused maps thus derived were merged into the Kir6.1_4_SUR2B_1_ maps to obtain composite maps for final modeling and structural analysis (Figs.1, S1).

To assess potential cooperativity of NBD dimerization of the four SUR2B subunits, we determined the number of particles with 2, 3, or 4 SUR2B subunits in the OD conformation by 3D classification of 410,772 symmetry expanded particles into 9 classes using masks that include Kir6.1 tetramer and either adjacent SUR2B or diagonal SUR2B subunits (Fig.S3). Of the 52,882 particles showing adjacent SUR2Bs in the OD conformation and 50,565 particles showing diagonal SUR2Bs in the OD conformation, 9,800 particles are common to both, which represent particles with 3 or 4 SUR2Bs in the OD state (Fig.S3). As these particle numbers are even lower than those calculated (∼78,000 for adjacent, ∼50,000 for opposite, and ∼20,000 for 3 and 4 SUR2Bs in the OD state) assuming independence and a probability of an NBD-dimerization event of ¼ (based on the fraction of particles showing OD conformation in symmetry expanded particles), we conclude there is no positive cooperativity for NBD dimerization under our experimental condition.

### Structural dynamics of SUR2B in vK_ATP_ channels

In the OD vK_ATP_ structure with dimerized NBDs, clear cryoEM densities for MgATP at the ATPase degenerate NBD1 site and MgADP at the ATPase consensus NBD2 site are observed. The TMD1, TMD2, NBD1 and NBD2 in this structure resemble the previously reported SUR2B-only structure bound to Mg-ATP/Mg-ADP and a K_ATP_ opener (PDB ID 7VLS)^23^. The IF vK_ATP_ structure where the two NBDs are clearly separated also shows bound Mg-nucleotide densities at both NBDs. However, the dynamic nature of NBD2 in the IF conformation limits resolution to allow definitive assignment of the density as MgATP or MgADP. Nonetheless, based on structures of pK_ATP_ or SUR proteins alone as well as other ABC transporters determined in the presence of MgATP/MgADP in the IF conformation, NBD2 is most likely bound to MgATP and is tentatively modeled as such. The IF conformation resembles the propeller-like conformation of vK_ATP_ structure we determined previously in the presence of ATP and the sulfonylurea inhibitor glibenclamide wherein ATP is bound at NBD1 and glibenclamide bound in a transmembrane pocket above NBD1 (more in Discussion)^14^.

A significant number of particles appear to be intermediate between the IF and OD states, with partially separated NBDs. To gain further insights, we performed 3D Variability Analysis (3DVA) in cryoSPARC^22^. The top seven principal components (eigenvectors) contributing to the variability within a C4-symmetry expanded set of particles combining the OD, IF and intermediate classes (Movie 1), of the OD state only (Movie 2) or of the IF state only (Movie 3) were analyzed (see Methods). Within well-resolved particles the top eigenvector is the NBD-dimerization to NBD-separation motion followed by the eigenvector capturing a motion of the SUR2B-NBDs coming closer or further away from Kir6.1-CTD due to a small rigid body movement of the SUR2B-ABC core (Fig.2, Movie 1). This rigid body motion is akin to that observed between propeller-like conformations previously reported in vK_ATP_ structures bound to inhibitory ATP and glibenclamide (Fig.S4)^14^, suggesting it may be an intrinsic property of the SUR2B-ABC core. Other variabilities appear to represent changes in the IDRs of SUR2B, including the 600-a.a. loop corresponding to the T1-N1 linker and the disordered Kir6.1-KCt. For variability within the OD and the IF classes, the rigid body movement of the SUR2B-ABC core resulting in the NBD1 being closer or further away from Kir6.1 is seen in both (Movies 2, 3). Additional variability within the OD class of particles includes a slight expansion and contraction of the extended Kir6.1-CTD as well as changes in density that may represent changes in the IDRs (Movie 2). In the IF class of particles, a major variability that stands out from the OD conformation is the disappearance of NBD2 density, consistent with lower resolution and increased dynamics of the NBD2 in the IF conformation (Movie 3).

**Figure 2.**
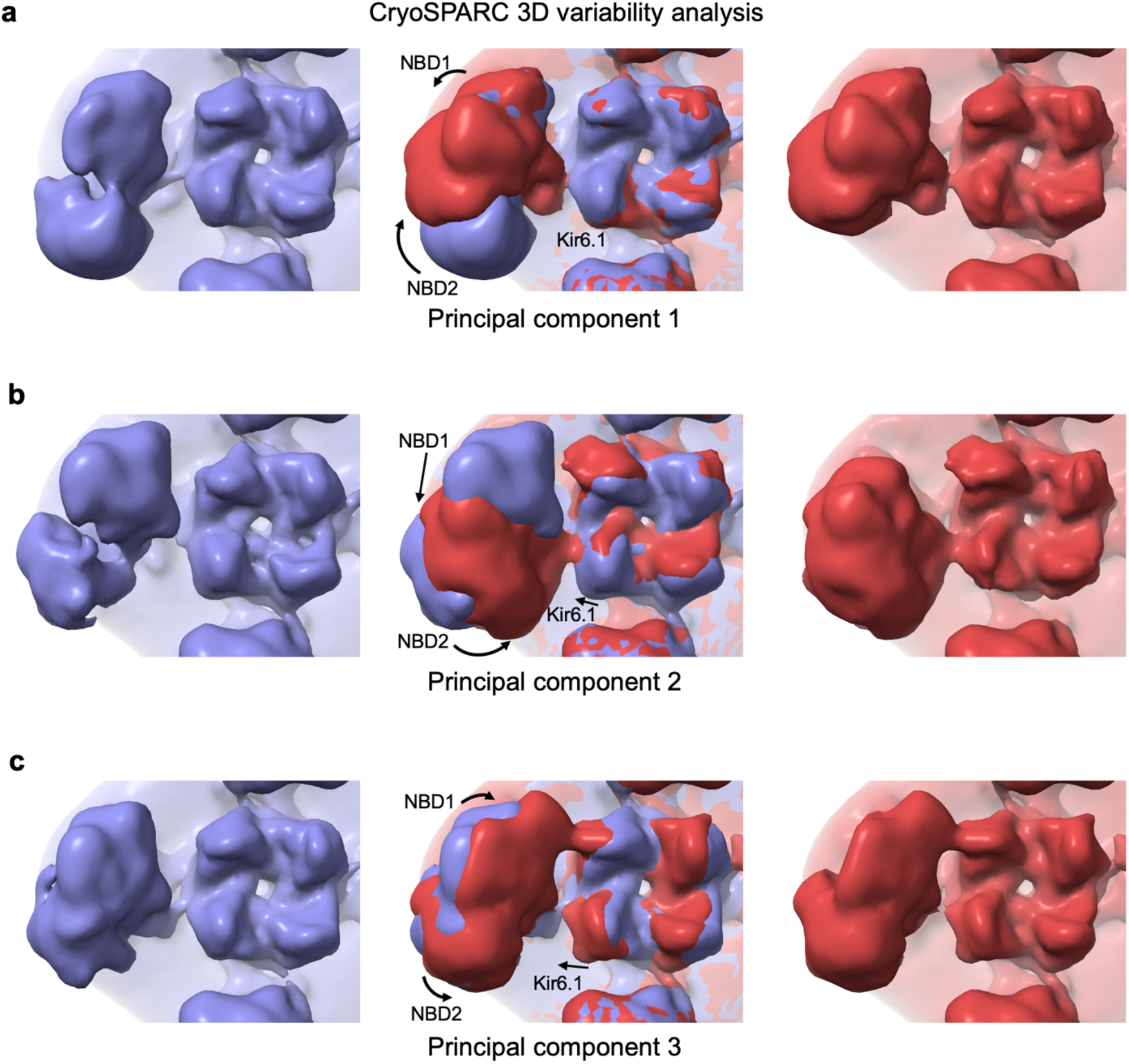
Dynamic motions of vascular K_ATP_ channel particles revealed by CryoSPARC 3D variability analysis. **(a)** A view from the cytoplasm of the conformational variability observed in principal component 1. The blue and red cryoEM maps represent the extreme ends of the conformational variability, with arrows showing the approximate direction and magnitude of movement from the blue map to the red map for each of the NBDs of SUR2B (NBD1, NBD2) and the Kir6.1 core. **(b)** The conformational variability observed in principal component 2 is shown in the same way as (a), showing a motion from an inward facing (IF) NBD-separated conformation to an occluded (OD) NBD-dimerized conformation. **(c)** The maps of the extreme ends of principal component 3 are shown in the same way as in (a) with the SUR2B-NBDs and Kir6.1-CTD tilting closer together.

Previously we reported two vK_ATP_ conformations bound to glibenclamide and Mg^2+^-free ATP: propeller (P)-like conformation, and quatrefoil (Q)-like conformation, both with SUR2B’s NBDs separated^14^. The Q-like structure differs from the P-like structure by a major counterclockwise rotation of the SUR2B-ABC core toward the Kir6.1 tetramer (Fig.S4). In the current study conducted in the presence of MgATP/MgADP, no Q-like conformation was observed (Fig.S4). Since the glibenclamide and Mg^2+^-free ATP cryoEM imaging condition in our previous study is non-physiological, the Q-like conformation may exist only in specific non-physiological conditions where the conformational freedom of the SUR2B-ABC module may be less constrained.

### Kir6.1 channel in closed conformation

In contrast to the dynamic SUR2B-ABC core, the structure of the Kir6.1 and SUR2B-TMD0 tetrameric core is stable and indistinguishable between the IF and OD conformations. In both, the Kir6.1 channel pore is closed as indicated by constrictions in the two cytoplasmic gates: the helix bundle crossing (F178) and the G-loop (G304, I305) (Fig.3). ATP is bound to all four inhibitory sites of the Kir6.1 tetramer (Fig.1). The C-linker, which connects the transmembrane helix M2 and the C-terminal cytoplasmic domain, is unwound into a loop stretching toward the cytoplasm such that the Kir6.1-CTD is in an extended conformation away from the membrane. This same extended conformation was previously reported for Kir6.1 in vK_ATP_ bound to glibenclamide and ATP where SUR2B was in NBD-separated IF conformation (Fig.3)^14^.

**Figure 3.**
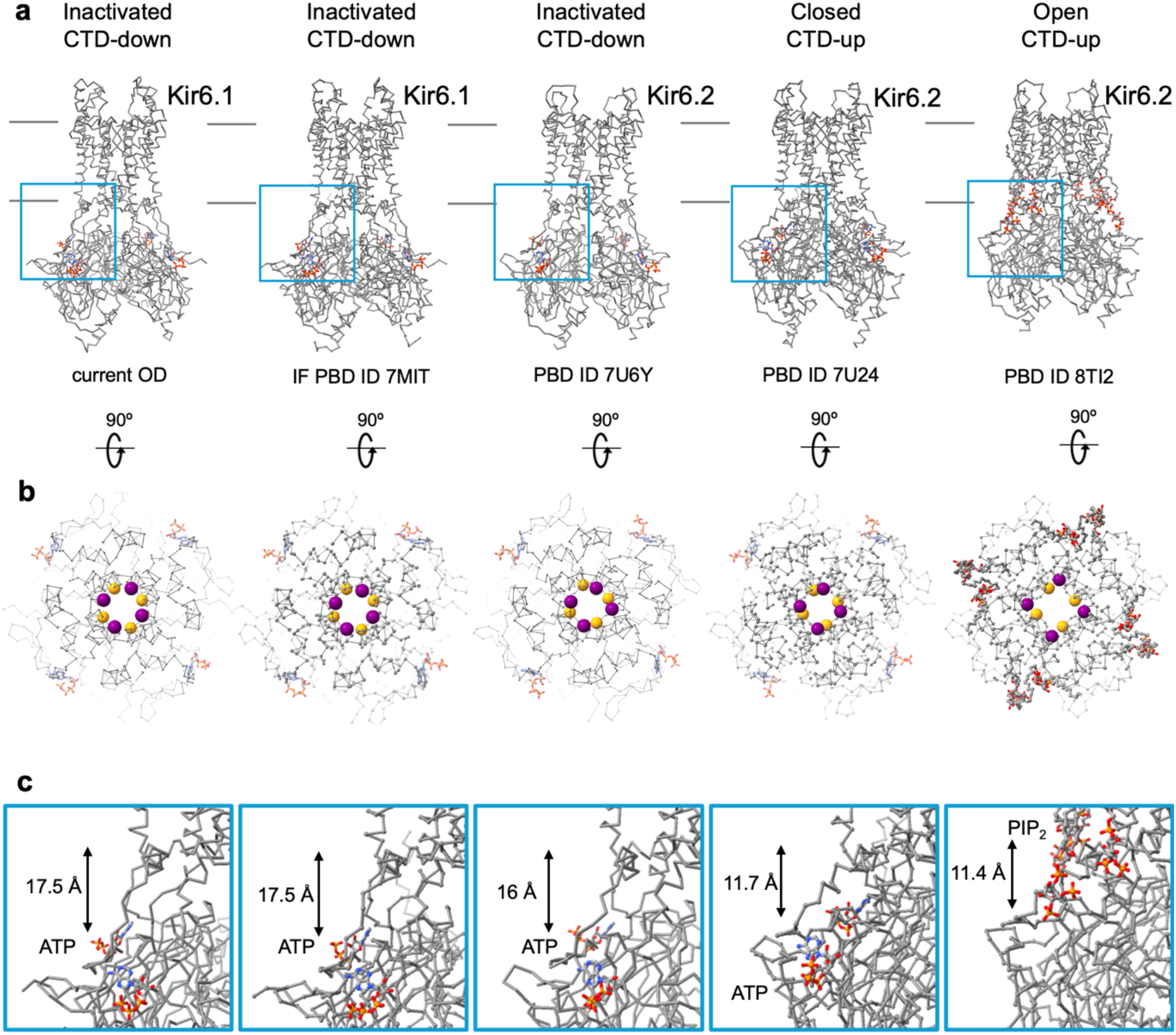
Comparison of Kir6.1 and Kir6.2 conformations. **(a)** CryoEM structures of Kir6.x shown as an alpha carbon (Cα) chain trace from the side with the position of the membrane marked (parallel grey lines) and with the zoomed in area shown in panel (c) indicated as a blue box. ATP or PIP_2_ are shown as sticks with atoms colored by element. From left to right: cryoEM structures of ATP-bound Kir6.1 in a CTD-down conformation (untethered from the membrane) corresponding to an inactive channel with SUR2B in a NBD-dimerized conformation; ATP-bound Kir6.1 in the CTD-down conformation corresponding to an inactive channel in the NBD-separated conformation (PDB ID 7MIT); ATP-bound Kir6.2 in the CTD-down conformation corresponding to an inactive channel in the ATP and glibenclamide (GBC)-bound NBD-separated conformation (PDB ID 7U6Y); ATP-bound Kir6.2 in the CTD-up conformation corresponding to a closed channel in the ATP and GBC-bound NBD-separated conformation (PDB ID 7U24); PIP_2_-bound Kir6.2 in the CTD-up conformation corresponding to an open channel (PDB ID 8TI2). **(b)** A rotated view of the same Kir6.x structures looking from the outside of the cell down the potassium pore, with the Cα of the key gating phenylalanine at the helix bundle crossing shown as a purple sphere (Kir6.1-F178, Kir6.2-F168), and the Cα of glycine at the G-loop gate shown as an orange sphere (Kir6.1-G304, Kir6.2-G295). **(c)** Zoomed-in side views of the area shown as a blue box in (a) focusing on the inner membrane interface with the distance shown between the helix bundle crossing and the G-loop gate.

In Kir channels the CTD is known to adopt two conformations, one tethered to the membrane conducive to interaction with PIP_2_ for channel opening, the other extended away from the membrane and counterclockwise rotated (sideview) relative to the tethered conformation representing an inactivated, closed, state^24–27^. The exclusively extended Kir6.1 configuration contrasts that observed for Kir6.2 bound to ATP in pK_ATP_ structures (Fig.3)^14,26,28^. In pK_ATP_, the majority of Kir6.2-CTD bound to ATP regardless of SUR1 conformation adopts the tethered position with a minor fraction in the extended position^26,27^. In contrast, in apo pK_ATP_ structure without ATP, Kir6.2-CTD is only in the extended position^26^ (Fig.3). That Kir6.1-CTD is found only in the extended position even when bound to ATP suggests Kir6.1-CTD is more resistant to rotate and move upward to the tethered position or that Kir6.1-CTD can move to the tethered position but is not stable enough in that position to be captured by cryoEM.

Unlike Kir6.2-containing K_ATP_ channels which are predominantly open in ATP-free solutions, Kir6.1-containing K_ATP_ channels are closed and require Mg-nucleotides for activity^3,29,30^, potentially reflecting a need for Mg-nucleotide induced SUR NBD dimerization to overcome the energy barrier for Kir6.1 to move its CTD to the membrane-tethered position. Analysis of particles with 1-4 SUR2B subunits in NBD dimerized states in our sample suggests there is no positive cooperativity among the SUR2B subunits for NBD dimerization and gives no evidence of upward movement of the Kir6.1-CTD even with multiple SUR2B subunits in the NBD-dimerized conformation. It is possible that ATP unbinding from Kir6.1 and SUR2B NBD dimerization are both needed for Kir6.1-CTD to move to the membrane-tethered position and that the ATP concentration present in our sample was too high for ATP dissociation from Kir6.1. Of note, all open pK_ATP_ channel structures to date employed mutations that eliminate inhibitory ATP binding at Kir6.2 or mutations that stabilized the channel in PIP_2_-bound open state^16,28,31^. Similar strategies may be necessary to understand how Mg-nucleotides open Kir6.1-containing vK_ATP_ channels.

### Intrinsically disordered regions

In K_ATP_ cryoEM structures reported to date, several regions are unresolved or only partially resolved under specific experimental conditions (reviewed in^12^), indicating they are highly dynamic^32,33^. Major K_ATP_ IDRs include the KNt (the N-terminal 1-32 residues of Kir6.1), the KCt (residues 365-424 residues of the Kir6.1 C-terminal), the T1-N1 linker (residues 618-664 between TMD1 and NBD1 of SUR2B; also referred to as the 600-loop), and the N1-T2 linker (residues 910-961 connecting NBD1 and TMD2 of SUR2B) (Fig.1). Strikingly, in both the IF and OD cryoEM maps from our sample, densities corresponding to Kir6.1-KNt and SUR2B N1-T2 linker (Fig.4) as well as weak and discontinuous densities for the SUR2B T1-N1 linker and Kir6.1-KCt are all observed (Fig.S5). Interestingly, the two best resolved IDRs—the Kir6.1-KNt and SUR2B N1-T2 linker—show distinct configurations and interactions in the IF structure versus the OD structure (Fig.4, Movie 4), suggesting they may have state-dependent structural and functional roles.

**Figure 4.**
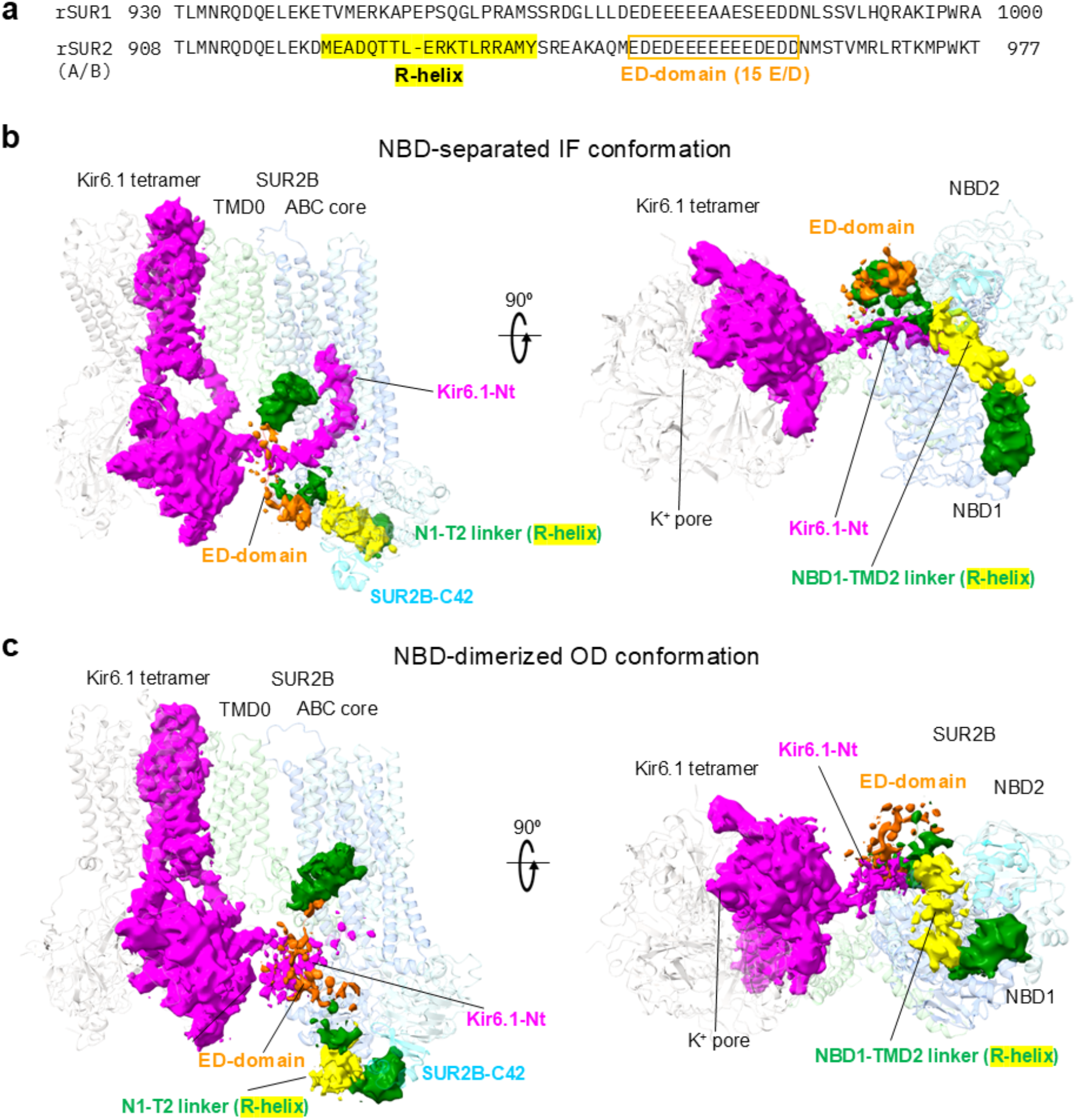
CryoEM maps of the SUR2B NBD1-TMD2 (N1-T2) linker including the R-helix and ED-domain and one Kir6.1 subunit of the vascular K_ATP_ channel. **(a)** Sequence alignment between SUR1 and SUR2(A/B) at the N1-T2 linker region showing that the R-helix region has three Prolines in SUR1, which disrupt alpha-helix formation, and that SUR1 has three neutral residues in the ED domain. **(b)** NBD-separated cryoEM map regions are shown at 2.5 rmsd (0.045 V), colored zones are shown within 5.0 Å of defined regions of the NBD-separated (IF) model. SUR2B residues 922-938 make up the R-helix portion of the NBD1 to TMD2 linker (N1-T2 linker) and are colored yellow. SUR2B residues 947-961 make up the ED-domain of the NBD1 to TMD2 linker (N1-T2 linker) and are colored orange, with multiple Glu and Asp residues able to interact with Arg and Lys residues of the KNt (magenta). The view from the side (*left*) and a rotated view from the cytoplasm (right) are shown. Note the non-continuous density of some of the IDRs at this contour indicates that these IDRs are highly dynamic. **(c)** NBD-dimerized (OD conformation) map regions are shown at 2.5 rmsd (0.048 V), colored zones are shown within 5.0 Å of the defined regions of the NBD-dimerized model, like in (b).

#### The Kir6.1 N-terminal disordered region (KNt)

In our SUR2B IF structure, the KNt extends into the cleft between the two TM bundles of SUR2B’s ABC core (Fig.4), as in previously reported closed pK_ATP_ and vK_ATP_ structures bound to inhibitory ATP and glibenclamide^14,34^. However, in pK_ATP_ channel structures where Kir6.2 is open, whether SUR1 NBDs are separated (such as in the SUR1/Kir6.2 open structure bound to PIP_2_)^28^ or dimerized (in the presence of MgATP/MgADP)^31^, the Kir6.2-KNt cryoEM density is absent. This suggests that KNt withdraws from the SUR-ABC core cavity when the Kir6 pore opens, but that SUR NBD-dimerization is not required to expel KNt from the cavity^28^. However, where KNt is when it is not in the SUR-ABC core cavity is unknown, as no clear KNt density was resolved in any of the published open pK_ATP_ structures^16,28,31^. Interestingly, here we observed amorphous cryoEM density for Kir6.1-KNt in our SUR2B OD state vK_ATP_ structure (Figs.4, S5). The KNt density is excluded from the SUR2B-ABC core cavity and is instead near the entrance of the cavity (Figs.4, S5), likely due to a more constricted cavity following NBD dimerization that can no longer accommodate the KNt.

#### The SUR2B N1-T2 linker

In both the IF and OD vK_ATP_ structures, cryoEM density for the SUR2B N1-T2 linker is apparent, albeit at lower resolution than the well-structured TMDs (Figs.4, S5). The SUR2B N1-T2 IDR, extending from residue 910-960, contains two sequences that have been implicated in functional regulation of K_ATP_ channels: one is a short α-helical structure called the R(Regulatory)-helix (residues 922-938)^23^ and the other is a stretch of 15 consecutive glutamate and aspartate residues designated the ED-domain (residues 947-960) (Fig.4)^33^.

The R-helix in the IF structure reported here sits between SUR2B’s two NBDs, akin to that observed previously in SUR2A and SUR2B single subunit-only IF structures^23^, and consistent with its proposed role of physically blocking NBD dimerization, hence channel activation. Interestingly, in our OD structure with SUR2B’s NBDs dimerized, the R-helix can no longer fit in the space between the NBDs and is found instead outside the dimerized NBDs (Figs.4, S5). Specifically, in contrast to a previously reported NBD-dimerized OD state structure of SUR2B without Kir (bound to MgADP/MgATP plus the channel activator levcromakalim or P1075) in which cryoEM density for the N1-T2 linker past residue 912 is missing^23^, our full vK_ATP_ structure with SUR2B in the OD conformation map clearly shows the N1-T2 linker extending as a helix for an additional 20 residues and then bending to follow along the groove of the dimerized NBDs where it interacts with the C42 module that is uniquely spliced in SUR2B (Figs.1, 4, S5).

C-terminal to the R-helix is the ED domain (Fig.4). Deletion or charge neutralization of this domain interrupts the SUR2A/Kir6.2 (cK_ATP_) response to MgADP, the pharmacological activator pinacidil, and the inhibitor glibenclamide^35^. In the vK_ATP_ IF structure the ED domain is in contact with the middle to C-terminal end of the KNt, which is inserted in SUR2B’s ABC core cavity (Fig.4). In the vK_ATP_ OD state structure, the density of the ED domain is in touch with an amorphous density corresponding to Kir6.1 KNt near the entrance of the SUR2B ABC core cleft (Fig.4). The resolution of the cryoEM density for both KNt and the ED domain is insufficient for sidechain modeling, suggesting they remain partially disordered or dynamic. Nonetheless, main chain models are possible. The structural interactions of the ED domain and the KNt in both the IF and OD conformations are consistent with the ED domain having a role in channel gating.

### MD simulations reveal interactions between vK_ATP_ IDRs

Because the cryoEM densities represent averaged conformations of the IDRs, we employed all-atom molecular dynamics (MD) simulations to further probe their conformational dynamics and interaction networks. Two sets of all atom simulations of the full SUR2B/Kir6.1 channel were performed using the IF or OD conformations as starting points (see Methods) each with five replicates of 500 ns duration.

To characterize interactions between individual IDRs, we first quantified persistent fragment-fragment contacts throughout the simulations. The analysis included the Kir6.1 KNt and KCt as well as the SUR2B T1-N1 linker (600-loop), the NBD1 700a.a.-loop, and the N1-T2 linker. Persistent interactions were observed both within individual SUR2B-Kir6.1 pairs and between neighboring channel subunits. The most prominent interaction in both conformations involved the N1-T2 linker and the Kir6.1 KNt (Figs.5, S6), suggesting that these two IDRs remain in close contact despite extensive conformational fluctuations within each IDR (Fig.S6). We therefore quantified residue-residue contact frequencies within this interface, focusing on KNt and the N1-T2 linker around the ED domain in both the IF and OD conformations. The corresponding residue interaction networks for both conformations are shown (Fig.5). Only residues with a minimum contact duration of 5% of the entire simulation time are included. Rather than being maintained by a few persistent residue pairs, the KNt-ED interface is stabilized by a continuously reorganizing network of transient residue-residue contacts.

**Figure 5.**
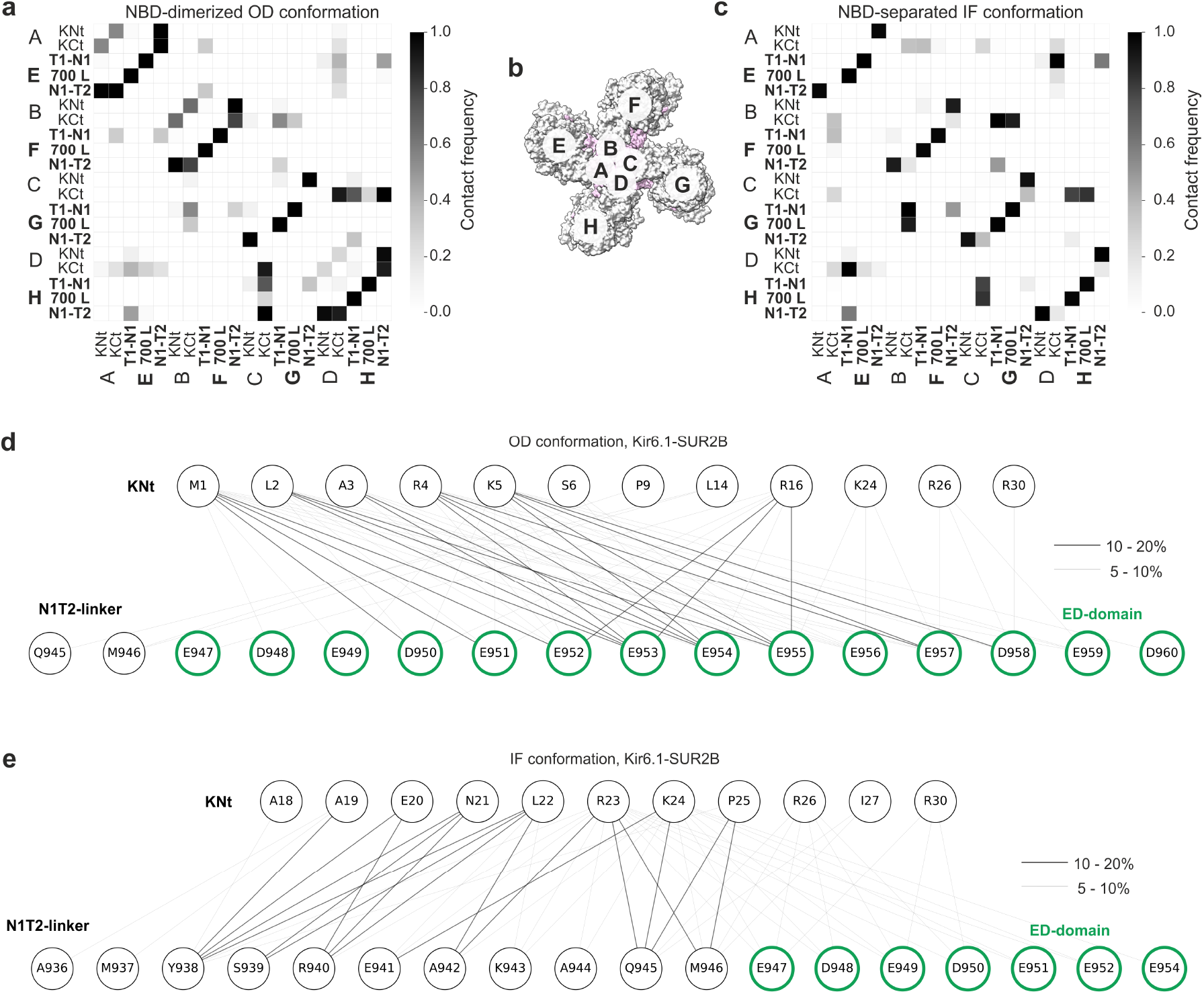
Fragment-based close-contact analysis of intrinsically disordered regions. **(a)** Heatmap showing the frequencies of close contacts between selected structural fragments in the NBD-dimerized (OD) conformation. **(b)** Schematic representation of the analyzed fragments. **(c)** Frequencies of close contacts in the NBD-separated (IF) conformation. **(d,e)** Contact networks between the SUR2B N1–T2 linker (ED residues circled in green) and the Kir6.1 N-terminus (KNt) in the NBD-dimerized (OD) conformation, where the KNt remains outside the SUR2B ABC core, and the NBD-separated (IF) conformation, where the KNt is inserted into the SUR2B ABC core. Edge thickness represents different contact-frequency ranges.

In the IF structure simulations, the N-terminal portion of the ED domain interacted with the middle to C-terminal portion of the KNt (such as R23, K24, and R26) as the distal N-terminal tail of Kir6.1 was inserted deeply in the SUR2B-ABC core cleft. In addition to the ED domain itself, frequent contacts were also observed with residues immediately preceding the ED domain within the N1-T2 linker (Figs.5, 6, S6), indicating that the interaction interface extends beyond the canonical ED-domain boundaries. In contrast, in the OD structure simulations, the interactions between KNt and ED-domain were primarily mediated by R4 and K5 from the distal part of KNt, and R16 from the middle part of KNt (Figs.5, S6). Intriguingly, these inter-fragment interactions were highly transient; almost every individual interaction lasted for no more than 20% of the total simulation time. Despite these differences in residue preferences, individual residue-residue contacts were highly transient, with nearly all interactions persisting for less than 10% of the cumulative simulation time. Nevertheless, the KNt and ED-domain remained associated throughout the simulations in both conformations, indicating that interface stability arises from continual exchange of transient interaction partners rather than persistent residue pairs (Figs.5, S6). This persistent fragment-level association is further illustrated by tracking centers of mass (Cα) of the ED domain (SUR2B residues 947-961) and three consecutive KNt segments (residues 1-7, 10-19, and 20-30), which remained spatially associated throughout the simulations despite continual rearrangement of residue-level contacts (Fig.S6).

In addition to its interactions with Kir6.1-KNt, SUR2B N1-T2 also made frequent contact with KCt of the same Kir6.1 subunit (Fig. 6). Moreover, the N1-T2 linker interacted with the neighboring SUR2B T1-N1 linker (600-loop), while the Kir6.1-KCt extended toward both the 600-loop and the NBD1 700-loop of the adjacent SUR2B subunit (Figs.6, S6, S7).

**Figure 6.**
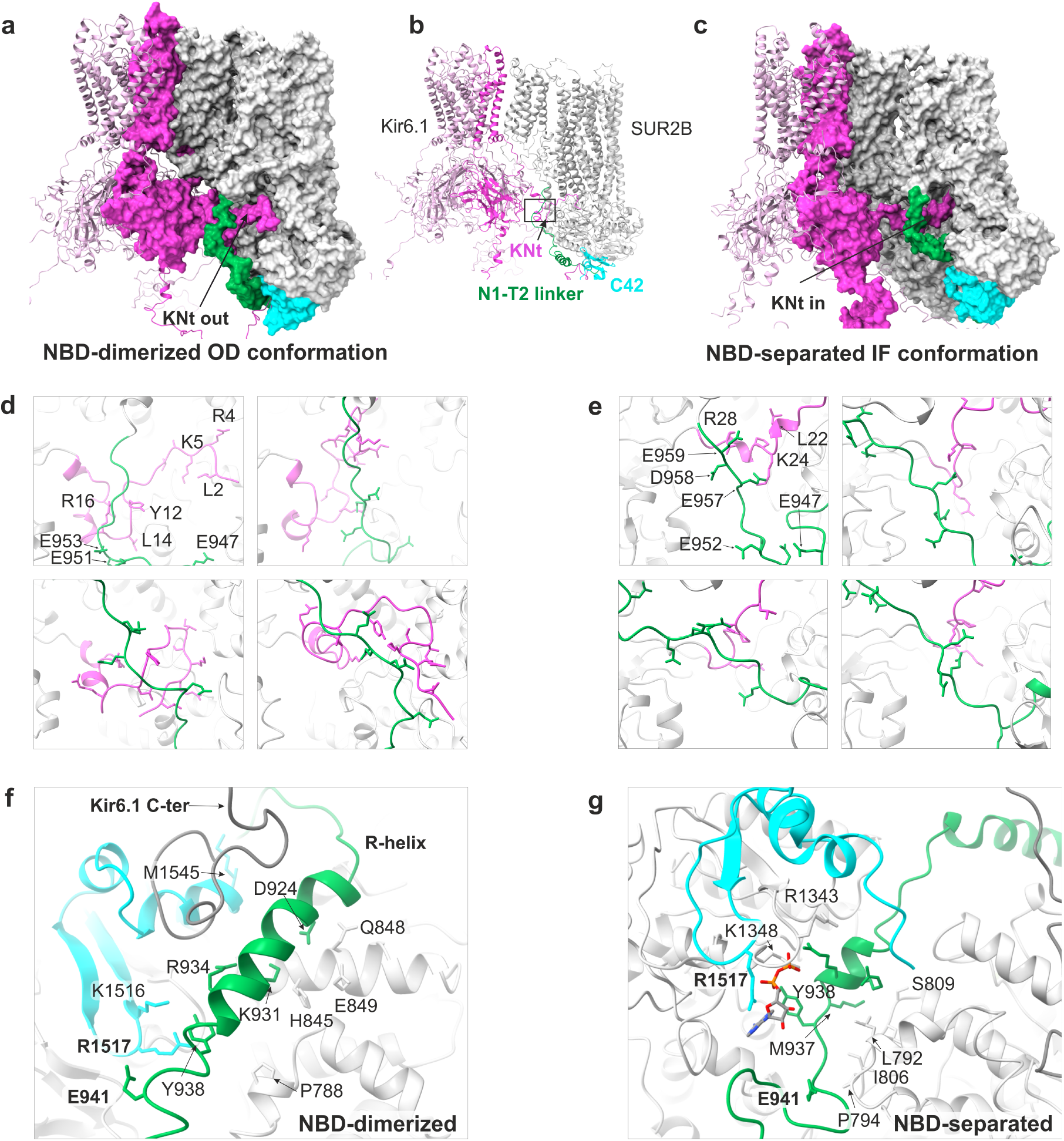
Representative molecular dynamics models and snapshots illustrating interactions between IDRs in vascular K_ATP_ channels. **(a)** Representative model of the NBD-dimerized (OD) conformation. **(b)** Overview of the analyzed intrinsically disordered regions (IDRs) with the boxed area indicating the region shown in panels d–g. **(c)** Representative model of the NBD-separated (IF) conformation. **(d,e)** Representative MD snapshots showing interactions between the Kir6.1 N-terminus (KNt) and the SUR2B ED domain in the NBD-dimerized and NBD-separated conformations, respectively. **(f,g)** Representative interactions of the SUR2B R-helix with neighboring structural elements in the NBD-dimerized and NBD-separated conformations.

Another notable interaction is between the SUR2B R-helix and C42 that is present in the OD structure (Fig.6). Unlike the KNt-ED interface, this interaction remained stable throughout all five 500 ns simulation replicates. Although individual residue-residue contacts were still transient, E941 within the R-helix consistently showed the highest interaction frequency with R1517 in C42 (Fig.6). These interactions may help to stabilize the SUR2B in the MgADP-stimulated OD conformation.

Together, these observations indicate that the cytoplasmic IDRs do not function as isolated flexible segments but instead form an interconnected interaction network spanning both intra-and inter-subunit interfaces.

## DISCUSSION

The study here presents the cryoEM structures of vK_ATP_ channels obtained in the presence of MgATP and MgADP. Significantly, these structures provide insights into the elusive IDRs, particularly the Kir6.1-KNt and the SUR2B N1-T2 linker, in both the SUR2B NBD-separated IF and NBD-dimerized OD conformations. Together with all-atom MD simulations of the full channel, they reveal significant and dynamic rearrangements of the KNt and N1-T2 linker between the IF and OD conformations, wherein the R-helix and ED domain in the N1-T2 linker engage with distinct regions in the Kir6.1 KNt and SUR2B NBDs to stabilize each conformation. The visualization of Mg-nucleotide dependent rearrangement of the K_ATP_ IDRs and evidence for the functional relevance offer novel insights into the structural and potential functional roles of these malleable elements.

### IDRs in MgADP/MgATP-dependent conformational switch in K_ATP_ channels

KNt was the first IDR in K_ATP_ channels to be visualized by cryoEM^26,34,36,37^. In both pK_ATP_ and vK_ATP_ IF structures bound to glibenclamide, KNt is inserted within the central cleft of SUR’s ABC-core adjacent glibenclamide, which stabilizes the KNt-SUR interface^12,14,26^. In pK_ATP_ structures obtained in MgATP/MgADP where SUR1 is in the NBD-dimerized OD conformation, the KNt cryoEM density is absent from the cleft due to constriction of the space between the two SUR1 TM bundles^31^. However, the whereabouts of the KNt in SUR1 NBD dimerized conformation remained a mystery. The vK_ATP_ isoform structures in this study provide a first glimpse of the movement of the KNt between the NBD-separated IF and NBD-dimerized OD conformations. In the vK_ATP_ IF conformation, the KNt sits in the SUR2B-ABC core cleft with its middle segment interacting with the ED domain. In the vK_ATP_ OD structure, the KNt is excluded from the central cleft and relocates to near the entrance of the cleft where its distal segment is held by the ED domain (Movie 4). This location affords Kir6-CTD the freedom to rotate to the open conformation. MD simulations show that multiple Glu/Asp residues from the ED domain interact with Arg/Lys residues of proximal KNt in the IF state and distal KNt in the OD state (Figs.5, S6). These interactions are transient but frequent, which may allow the ED domain to stabilize the KNt in two distinct positions during SUR2B transition between IF and OD conformations without locking KNt in specific positions. In our previous MD study of pK_ATP_, we proposed that pK_ATP_ IDRs act as structural links mediating communication between different channel domains^32^. The MD simulations of vK_ATP_ in the present study based on cryoEM structures of both the IF and OD conformations reveal distinct interaction networks adopted by the vK_ATP_ IDRs in the two functional states, thus providing direct support for this concept. The structures also directly demonstrate that the KNt moves in and out of the SUR ABC core cleft during MgATP/MgADP-induced SUR IF and OD conformational switch, highlighting the importance of the KNt in physiological regulation of K_ATP_ channels.

The R-helix in the N1-T2 linker may also serve double duty. First, the R-helix wedges between the two NBDs when MgATP/MgADP concentrations do not favor NBD dimerization^23^. Second, when the condition favors NBD dimerization, the R-helix moves outside the dimerized NBDs and interacts along the groove of the dimerized NBDs with SUR2B-C42, which may help stabilize the NBDs in the dimerized conformation and enhance MgADP stimulation of the channel. However, we emphasize that the proposed functional roles are speculative as the lack of detectable activity of SUR2B/Kir6.1 channels in nucleotide-free solutions in isolated membranes^38–40^ precludes functional testing of these hypotheses.

Notable sequence variations exist in the IDRs of different SUR and Kir6 isoforms^12^. SUR1 lacks the R-helix and its ED-domain corresponding sequence is broken up by non-ED residues (Fig.4). The C42 sequence of SUR1 is also different from SUR2A and SUR2B, although much more homologous to SUR2B than SUR2A. These differences likely explain why no R-helix density or N1-T2 density were observed in pK_ATP_ structures published to date. Our previous MD simulations of SUR1 suggested that the N1-T2 linker is capable of sampling conformations in which it transiently occupies the space between the NBDs, despite lacking persistent helical structure^26,32^. The current cryoEM structures and previous structures of SUR2A or SUR2B protein alone show that in SUR2A and SUR2B this region adopts a defined R-helix to regulate NBD organization in a more ordered manner^23^. Between Kir6.1 and Kir6.2, there is high conservation in KNt, but Kir6.2-KCt is 24 residues longer. These sequence variations may underlie the different IDR dynamics in the different K_ATP_ channel proteins as reflected by whether the various IDRs are captured by cryoEM. Consistent with this idea, our MD simulations show that the Kir6.1- and Kir6.2-KCt and neighboring disordered segments explore markedly different conformational space in vK_ATP_ (Fig.S7) and pK_ATP_^32^. Such differences may contribute at least partially to the difference in the nucleotide sensitivity of the different K_ATP_ isoforms.

### Versatility of the N1-T2 linker in ABC transporters

The N1-T2 linker IDR has been captured by cryoEM and implicated in functional regulation of several other type IV ABC transporters, including the Cl^-^ channel cystic fibrosis transmembrane conductance regulator (CFTR)^41^, the multidrug resistance protein MRP2^42,43^, and the yeast transporter Ycf1 (Fig.7)^44–46^. For example, in CFTR and Ycf1, the N1-T2 linker, which contains a R-domain with multiple phosphorylation sites, is seen lodged in the ABC core cleft when unphosphorylated to prevent CFTR or Ycf1 function but moves out of the ABC core cleft upon phosphorylation to allow nucleotide regulation of Cl^-^ conductance in CFTR or transport in Ycf1 (Fig.7). In MRP2, the N1-T2 linker has been proposed to play an autoinhibitory role by occupying the substrate binding site when the NBDs are separated, only becoming dislodged when substrates are at sufficiently high concentrations to allow transport.

**Figure 7.**
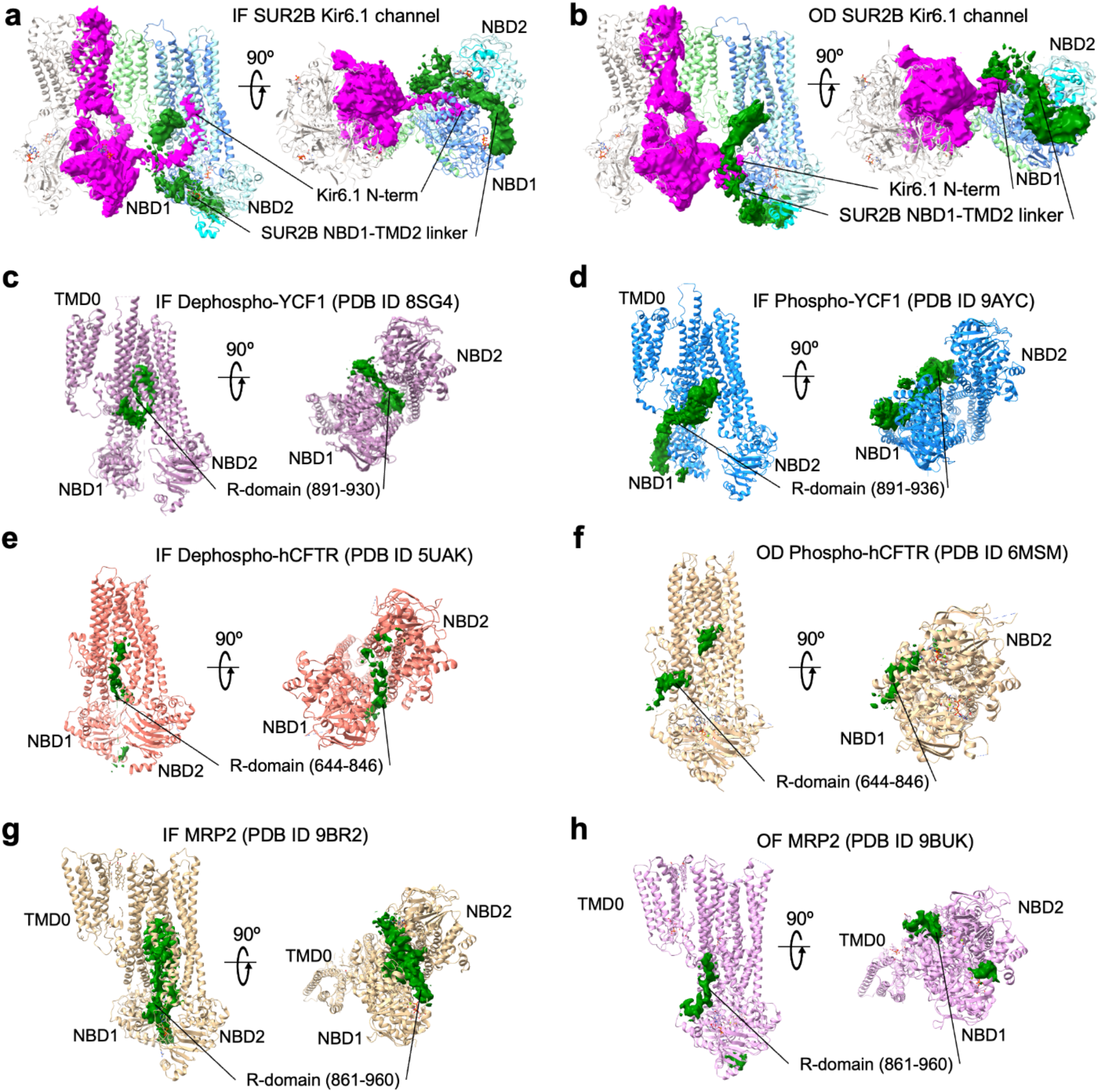
Comparison of the N1-T2 linker in type IV ABCC transporters. The cryoEM densities assigned to be a portion of the N1-T2 linker (also known as the R-domain in YCF1, CFTR, and MRP2) in all ABCC proteins shown here are colored green at low contours. **(a)** Vascular K_ATP_ channel reported here in a SUR2B NBD-separated inward-facing (IF) conformation with the cryoEM map density of SUR2B N1-T2 shown at 0.04 V contour (green). The cryoEM density of one Kir6.1 with its N-terminus (KNt) inside the ABC core central cleft of SUR2B is shown in magenta) at the same contour. **(b)** Vascular K_ATP_ channel reported here in a SUR2B NBD-dimerized occluded (OD) conformation showing KNt out of the ABC core near the N1-T2 linker. **(c)** Yeast YCF1 transporter in an IF conformation (PDB ID 8SG4) with map regions for a dephosphorylated R-domain shown at contour 0.03 V (EMD_40451). **(d)** Yeast YCF1 transporter in an IF conformation (PDB ID 9AYC) with its phosphorylated R-domain in the N1-T2 linker cryoEM density at contour 0.03 V (EMD_43985). **(e)** Human CFTR (hCFTR) in an IF conformation (PDB ID 5UAK) with map regions for a dephosphorylated R-domain shown at 1.4 rmsd (EMD_8516). **(f)** hCFTR in an OF conformation (PDB ID 6MSM) with map regions for a phosphorylated R-domain shown at 0.3 V contour (EMD_9230). (g) Multi-drug resistance protein 2 (MRP2) in an IF conformation (PDB ID 9BR2) with map regions for the R-domain shown at a contour of 0.04 V (EMD_44833). **(h)** MRP2 in an outward-facing NBD-dimerized (OF) conformation (PDB ID 9BUK) with map regions for the R-domain shown at 0.04 V (EMD_44911).

Compared to other type IV ABC transporters, SUR proteins are unique in not having any known transport or ion channel activity themselves but instead function as regulatory subunits of the K_ATP_ channel complex to modulate the activity of Kir6.1 or Kir6.2^47^. Here, SUR has acquired a tethered pseudosubtrate, namely the KNt of Kir6.1 or Kir6.2, which comes in and out of the ABC core central cleft of SUR to regulate the activity of the Kir6 channel, resembling the R-domain of CFTR and MRP2 (Fig.7). The SUR N1-T2 linker, at least for SUR2B shown here, rather than being regulated by phosphorylation to move in and out of the ABC core central cavity, as in other ABC transporters mentioned above, has evolved to harbor an ED domain that resembles a permanently phosphorylated R-domain to interact with the pseudosubstrate, KNt, facilitating channel gating regulation. Interestingly, in SUR1 the ED domain equivalent sequence contains several non-ED residues and the SUR1 N1-T2 linker has not been resolved in any of the pK_ATP_ cryoEM structures. It remains to be determined whether the SUR1 N1-T2 linker is subject to posttranslational modifications such as phosphorylation. Regardless, the examples discussed herein highlight the N1-T2 linker as a versatile IDR for regulating ABC protein structure and function.

In summary, our study of the vK_ATP_ channel illuminates how flexible IDRs such as the KNt and the N1-T2 linker regulate channel structure and activity through transient and ligand-dependent interactions with the stable domains and with one another. Pharmacological agents targeting such interactions, as exemplified by the anti-diabetic drugs sulfonylureas which stabilize KNt and SUR ABC core central cavity^26^, could be exploited for therapeutic development in the future.

## Methods

### Cell lines

COSm6 cells (RRID:CVCL_8561) were used for K_ATP_ channel expression. HEK-AD293 cells (RRID:CVCL_0063; from Agilant Technologies catalog number 240085) were used for production of recombinant adenovirouses encoding FLAG-SUR2B and Kir6.1, described previously^14^. Both COSm6 and HEK-AD293 cells were cultured in high-glucose DMEM medium (GIBCO) supplemented with 10% Fetal Bovine Serum (Fisher Scientific), 100 U/mL penicillin, and 100 U/mL streptomycin at 37°C with 5% CO_2_.

### Protein expression and purification

Genes encoding rat Kir6.1 and N-terminal FLAG-tagged (DYKDDDDK) SUR2B were cloned into pShuttle vectors and then the AdEasy vector (Stratagene), and packaged into recombinant adenoviruses, which were used for protein expression as described previously^14^. COSm6 cells grown to mid-log in 15 cm tissue-culture plates were infected with adenoviruses packaged with Kir6.1, SUR2B, and tTA, using multiplicity of infections (MOIs) optimized empirically. Note the pShuttle vector used for SUR2B contains a tetracycline-regulated response element, necessitating co-infection of a tTA (tetracycline-controlled transactivator) adenovirus for SUR2B expression. At ∼48 hours post-infection, cells were harvested by scraping and cell pellets were frozen and stored at −80°C until purification.

SUR2B/Kir6.1 channels were purified similar to previously described^48^, except after elution with a buffer containing 0.2 M NaCl, 0.1 M KCl, 0.05 M HEPES pH 7.5, 0.05% glyco-diosgenin (GDN), and 0.25 mg/mL FLAG peptide, eluant was loaded onto a Superose-6 increase 10/300 column at 0.5 mL/min using an AKTA FPLC (Cytiva®), and peak fractions of the fully assembled octameric channel were collected and concentrated using an Amicon® Ultra – 0.5mL Centrifugal Filter with Ultracel®-100,000 molecular weight cutoff^49^. Purified and concentrated SUR2B/Kir6.1 channels were used immediately for cryo-EM grid preparation.

### CryoEM sample preparation and data acquisition

To increase protein adsorption to the cryoEM grids, and also to mitigate selective-orientation of K_ATP_ channel particles that occurs on commercial carbon surfaces, graphene-oxide (GO) grids were prepared similar to previously described^14,50,51^. Briefly, Au Quantifoil R1.2/1.3 400 mesh grids (SPI®) were glow-discharged for 60 seconds at 15 mA with a Pelco EasyGlow®, and 4 µL of 1 mg/mL Polyethylenimine (PEI, 40,000 MW, Polyscienes Inc.) in 25 mM HEPES pH 7.9 was applied to each grid and incubated for 2 minutes followed by two washes with water. Then, 0.1 mg/mL GO was vortexed vigorously for at least 5 minutes and applied to the grid and incubated for 2 minutes followed by two washes with water, and used within 3 hours for sample vitrification.

To prepare cryoEM samples, 3.5 µL of purified K_ATP_ channel complex in a buffer containing 1 mM ATP, 1 mM ADP, 4 mM Mg^2+^ and 10 µM VU0542270 was loaded onto GO-coated grids. After 20 s at 6 °C with a humidity of 100%, the grids were blotted for 3 s with Horizontal and Vertical blot positions at 42.6 mm and 3.7 mm, respectively, and cryo-plunged into liquid ethane cooled by liquid nitrogen using a Leica EM GP2 Automatic Plunge Freezer in the Pacific Northwest Center for Cryo-EM (PNCC) that is jointly operated by Oregon Health & Sciences University (OHSU) and Pacific Northwest National Laboratory (PNNL).

Single-particle cryo-EM data was collected on a Titan Krios 300 kV cryo-electron microscope (ThermoFisher Scientific) in the PNCC, with a multi-shot strategy using beam shift to collect 27 movies per stage shift, assisted by the automated acquisition program SerialEM. Images were recorded on the Gatan BioContinuum K3 direct-electron detector in super-resolution mode, post-GIF (20eV window), at 105,000x magnification (calibrated image pixel-size of 0.825 Å); nominal defocus was varied between −0.5 and −2.5 µm across the dataset. The dose rate was kept around 14 e^-^/Å^2^/sec, with a frame rate of 18 frames/sec, and 70 frames in each movie (i.e. 3.9 sec exposure time/movie), which gave a total dose of approximately 55 e^-^/Å^2^. Two grids that were prepared in the same session using the same protein preparation were used for data collection, and from these two grids, 7,151 and 14,814 movies (21,965 movies total) were recorded (Fig.S1).

To prepare cryoEM samples in the absence of inhibitor, 3.5 µL of purified K_ATP_ channel complex in a buffer containing 1 mM ATP, 1 mM ADP, and 4 mM Mg^2+^ was loaded onto GO-coated grids. After 20 sec at 6°C with a humidity of 100%, the grids were blotted for 3 sec with Horizontal and Vertical blot positions at 42.6 mm and 3.7 mm, respectively, and cryo-plunged into liquid ethane cooled by liquid nitrogen using a Leica EM GP2 Automatic Plunge Freezer in the PNCC.

For samples in the absence of inhibitor, single-particle cryo-EM data was collected on a Titan Krios 300 kV cryo-electron microscope (ThermoFisher Scientific) in the PNCC, with a multi-shot strategy using beam shift to collect 27 movies per stage shift, assisted by the automated acquisition program SerialEM. Images were recorded on the Gatan BioContinuum K3 direct-electron detector in super-resolution mode, post-GIF (20eV window), at 81,000x magnification (calibrated image pixel-size of 1.059 Å); nominal defocus was varied between −0.5 and −2.5 µm across the dataset. The dose rate was kept around 15 e^-^/Å^2^/sec, with a frame rate of 18 frames/sec, and 70 frames in each movie (i.e. 3.9 sec exposure time/movie), which gave a total dose of approximately 55 e^-^/Å^2^, 11,575 movies were recorded (Fig.S2).

### CryoEM image processing

Super-resolution dose-fractionated movies were gain-normalized by inverting the gain reference in Y and rotating upside down, corrected for beam induced motion, aligned, and dose-compensated using Patch-Motion Correction in cryoSPARC2^20^ without binning. Parameters for the contrast transfer function (CTF) were estimated from the aligned frame sums using Patch-CTF Estimation in cryoSPARC2^20^ and binned by 2 with Fourier cropping. Micrographs were manually curated using sorting by curate exposures, resulting in 5554 and 12,948 micrographs from each of the two grids. The resulting 18,502 dose-weighted motion-corrected summed micrographs were used for subsequent cryoEM image processing. Particles were picked automatically using 2D templates based on 2D classes of around 1000 manual picked particles. Particle images were extracted based on template pick locations using a 512×512 pixel box sampled to 256×256 pixels and cleaned by three rounds of 2D classification in cryoSPARC2^20^. The particle stack after 2D classification rounds contained 102,693 particles, which were then re-extracted at 512×512 pixel box size from 16,087 micrographs that had good picks, and used for ab initio reconstruction without symmetry restraints imposed (Fig.S1, Table S1). Subsequent homogeneous refinement, using a mask of the K_ATP_ channel particle that included the micelle without symmetry restraints, of all 102,693 particles gave maps with 3.5 Å resolution reconstruction by GSFSC cutoff of 0.143^21^. Homogeneous refinement with C4 symmetry imposed but without a mask had 4.1 Å resolution reconstruction by GSFSC cutoff of 0.143 (Fig.S1)^21^. For data in the absence of inhibitor, movies were processed similarly and 11,575 micrographs were used for particle picking. After 2D classification, 32,785 particles from good 2D classes were used for homogenous refinement and gave maps with 6.3 Å resolution without symmetry restraints, and 4.4 Å resolution with C4 symmetry imposed (Fig.S2).

To identify structural conformations of the K_ATP_ channel within the particles, particles were 4-fold symmetry expanded to give a set of 410,772 particles, which were classified into six main classes using focused 3D classification in CryoSPARC with a mask focused on the NBDs of SUR2B^52^. One class had poorly resolved density for NBD2 that could represent damaged/dissociated particles on the grid, but which is likely due to a highly dynamic structure of NBD2, perhaps not bound to MgATP/MgADP. The missing NBD2 is similar to what we observe for the open pancreatic K_ATP_ channel that is in the absence of MgATP/MgADP^28^, and was previously reported for a 3D class of SUR2B-only channels in the presence of MgATP and repaglinide^23^. The other five classes of particles in two NBD-dimerized conformations, two NBD-separated conformations, or K_ATP_ channel particles with NBDs in an apparent intermediate state that is in-between IF and OD (Fig.S1).

Masks for classification and FSC calculation during refinement included the Kir6.1 tetramer plus one SUR2B subunit or focused on the NBDs were generated using molmap in ChimeraX^53^, with a threshold of 0.04 V applied and resampled on the full map grid (512×512×512 pixels). Volume tools in cryoSPARC were used to dilate the mask by 5 pixels and apply soft padding of 15 pixels^20^. For all cryo-EM maps, the mask-corrected Fourier shell correlation (FSC) curves were calculated in cryoSPARC2, and the resolutions were reported based on the 0.143 criterion^54^. A similar data processing strategy was used for data collected in the absence of VU270, which yielded lower-resolution OD and IF conformations (Fig.S2).

To gain insights into channel protein dynamics, we performed 3D Variability Analysis (3DVA) in cryoSPARC^22^. A C4-symmetry expanded set of particles that included particles from well-resolved classes, a set of particles from the two OD states, or a set of particles from the IF states were analyzed. The top seven principal components of variability (eigenvectors) contributing to the variability within those particles were analyzed and movies were made using 3D Variability Display in cryoSPARC^22^, and ChimeraX (movies 1, 2, 3)^53^.

To further explore the cooperation of NBD-dimerization for adjacent or opposite SUR2B subunits using our set of 410,772 C4-symmetry expanded vK_ATP_ channel particles, we examine if adjacent NBD-dimerized subunits are more likely to occur retaliative to opposite NBD-dimerized subunits, and compare that with what would be expected for theoretical probability distributions for independent NBD-dimerization events. We used a focused 3D-classification strategy in CryoSPARC^52^, with a mask covering two SUR2B subunits that were either adjacent, or opposite, each parsed into 9 classes (Fig.S3). For theoretical calculations we conservatively underestimated a probability of dimerization as *p* = 0.25 based on the number of particles with one subunit in the OD conformation (Fig.S1), and estimated the theoretical probability distribution assuming no-cooperation as independent Bernoulli trials with the probability of each subunit not experiencing dimerization as 0.75. For example, the probability of all SUR2B subunits not being NBD-dimerized was calculated as 0.75^4, and all being NBD-dimerized was calculated as 0.25^4.

### Model building

To create initial structural models, the structure of vascular K_ATP_ channel (PDB ID 7MIT) was fit into the reconstructed density for the full K_ATP_ channel particles in both the NBD-separated and the NBD-dimerized conformations using Chimera,^55^ and then refined in Phenix^56^ as separate rigid bodies corresponding to TMD (32-171) and CTD 172-352) of Kir6.1 and TMD0/L0 (1-284), TMD1 (285-614), NBD1 (615-928), NBD1-TMD2-linker (992-999), TMD2 (1000-1319) and NBD2 (1320-1582). The Kir6.1 tetramer transmembrane region and cytoplasmic domain fit well into both the NBD-separated and the NBD-dimerized cryoEM densities, with all structures of Kir6.1 having the cytoplasmic domain in an extended conformation down away from the membrane and bound to ATP.

For the NBD-separated conformation, generally the SUR2B model from PDB ID 7MIT fit well but there was additional density not fit by the model corresponding to a Mg-nucleotide bound to NBD2 of SUR2B, as well as density for the N1-T2 linker extending between the NBDs like the R-helix previously reported for SUR2A-only and SUR2B-only structures^23^. Compared with the glibenclamide-bound PDB ID 7MIT structure, there was not clear density for an inhibitor and alternative rotameric positions for sidechains around the inhibitor binding site were observed for residues R381, Q1164, Y1205, and R1263. Density for the N-terminus of Kir6.1 extended into the ABC-core of SUR2B, with changes in the model built manually using Coot^57^. The resulting model was further refined using *Coot* and *Phenix* iteratively until the statistics and fitting were satisfactory (Table S1). The model contains residues 1-366 for Kir6.1 chain A (missing C-terminal residues 367-424), with residues 1-30 not modeled for chains B, C, and D. The IF model contains residues 1-1545 for SUR2B except for two loop regions (619-664 and 734-740), and part of the N1-T2 linker after the R-helix (940-960). Loop regions that showed signs of disorder had most sidechains of residues in these regions stubbed at Cβ and manually refined into the map.

For the NBD-dimerized conformation, the TMD0 and the cytoplasmic linker L0 from the SUR2B model from PDB ID 7MIT fit the NBD-dimerized cryoEM maps well, with structural deviations associated with NBD-dimerization seen starting at the first TM helix in TMD1 (TM helix 6), which shifts as a ridged body with the cytoplasmic side more anchored and the extracellular portion shifting about 5 Å towards a more extended propeller conformation. The first TM helix of TMD2 (TM helix 12) shifts about 15 Å on the cytoplasmic side and about 12 Å on the extracellular region. A published model of SUR2B-only in the NBD-dimerized occluded conformation (PDB ID 7VLR)^23^ was used for comparisons when modeling the TMD1, NBD1, TMD2 and NBD2 regions of our vK_ATP_ channel in the Mg-nucleotide bound NBD-dimerized conformation. The OD model contains residues 1-366 for Kir6.1 chain A (missing C-terminal residues 367-424), with residues 1-30 not modeled for chains B,C, and D. The OD model contains residues 1-1545 for SUR2B except for an extracellular loop (1019-1027), and two loop regions (613-664 and 729-743).

In addition to protein density, two N-acetylglucosamine (NAG) molecules per SUR2B monomer, which are common cores of N-linked glycosylation, are modeled in the distinctively large density at the side chain of SUR2B-N9. ATP bound to the inhibitory site on Kir6.1-CTD and MgATP/MgADP were modeled in the SUR2B-NBDs. Ordered lipids were modeled in both conformations and one ordered GDN molecule was built in the OD model. The resulting model was further refined using *Coot*^57^ and *Phenix*^56,58,59^ iteratively until the statistics and fitting were satisfactory (Table S1). All structure figures were produced with UCSF Chimera^55^, ChimeraX^53^, and PyMol (http://www.pymol.org).

### MD simulations

The IF and OD cryoEM structures consisting of the Kir6.1 tetramer with one SUR2B subunit (4K1S) were used as starting models. In these models, the KNt of one of the Kir6.1 subunits (KNt) and the N1-T1 of SUR2B were present. Since intrinsically disordered regions (IDRs) interact significantly with neighboring domains, to obtain reliable information about their contacts, we proceeded with the full channel (4K4S) system. The Schrödinger software was used to expand the initial 4K1S model derived from the cryoEM map to 4K4S by copying the SUR2B protein and placing it in the same position relative to each Kir6.1 subunit as in the original Kir6.1-SUR2B assembly pair. Initial conformations of missing IDRs were generated using AlphaFold2^56^ combined with Afflecto^60^ predictions. The system also included ligands found in the original structure including the inhibitory ATP at Kir6.1 and MgATP/MgADP at the NBDs of SUR2B, all automatically protonated for pH 7.0. The system was initially minimized.

The system described above was initially minimized and then embedded in a 1-palmitoyl-2-oleoyl-phosphatidylcholine (POPC) lipid bilayer with water (TIP3) and 0.2 M KCl using CHARMM-GUI ^61,62^. The CHARMM36m force field was applied to the entire system^63^. Ligand parameters were generated automatically. After minimization, the system underwent preliminary equilibration, which included a series of 7 consecutive NVT and NPT simulations with gradually relaxed restraints on the system. Subsequently, while maintaining restraints (fc = 200 kJ/mol/nm²) on the backbone of the region with a known cryoEM structure, a 100 ns NVT simulation (5 independent replicates, starting from minimization) was conducted to allow the artificially added IDRs to equilibrate properly, losing information about their initial, somewhat arbitrarily chosen structure. This also provided sufficient time for the membrane to equilibrate based on previous simulations^32,49,64,65^. Note, it was important to ensure that the starting configuration of the added loops was different in each chain to sample the biggest possible conformational ensemble of IDRs in the production MD.

Production MD simulations were conducted for 500 ns, with 5 replicates, under NPT conditions. The pressure coupling was set to the C-rescale method, while the temperature coupling was performed using the v-rescale algorithm. The reference temperature was maintained at 310 K, and the time step for the simulation was set to 0.002 ps. Simulation data were saved every 50 ps. The simulations were performed using Gromacs 2023. Trajectories were aligned to the TMD helices of the Kir6.1 subunits.

### Data Availability

The coordinates of the structural models of the SUR2B/Kir6.1 vK_ATP_ channel and associated cryo-EM maps have been deposited in the Protein Data Bank (PDB) and the Electron Microscopy Data Bank (EMDB) under the following accession numbers: Kir6.1/SUR2B K_ATP_ channel in an NBD-separated inward-facing conformation in the presence of Mg Nucleotides (PDB ID 38LY/EMD-78915); Kir6.1/SUR2B K_ATP_ channel in an NBD-dimerized occluded conformation in the presence of Mg Nucleotides (PDB ID 38KK/EMD-78878). Additional cryoEM maps deposited include: SUR2B Kir6.1 K_ATP_ channel in NBD-separated inward-facing conformation, consensus refinement of Kir6.1 tetramer plus one SUR2B subunit (EMD-78917); SUR2B Kir6.1 K_ATP_ channel in NBD-separated inward-facing conformation, focus on one SUR2B subunit (EMD-78916); SUR2B/Kir6.1 K_ATP_ channel in an NBD-dimerized occluded conformation, consensus refinement of Kir6.1 tetramer plus one SUR2B subunit (EMD-78876); SUR2B Kir6.1 K_ATP_ channel in NBD-dimerized occluded conformation, focused refinement on one SUR2B subunit (EMD-78877); SUR2B/Kir6.1 K_ATP_ channel in the presence of MgATP/MgADP only in an inward-facing NBD-separated conformation (EMD-78802); SUR2B/Kir6.1 K_ATP_ channel in the presence of MgATP/MgADP only in the occluded NBD-dimerized conformation (EMD-78801).

All data and MD input files required to reproduce the numerical results reported in this paper are available in: https://urldefense.com/v3/_https://doi.org/10.18150/JBA3SJ_;!!Mi0JBg!OkQDv6645vp6Vmyk7zASKdTSzMNviZa0YSWsD-NmtncKk57UJ2SPKXL7QuW5YdNKd0Oy3 q3RksIDIg$

## Supporting information

Movie 1. 3D variability for all well-resolved particles

Movie 2. 3D variability for OD particles

Movie 3. 3D variability for IF particles

Movie 4. vKATP IF conformation to OD conformation transition

Molecular dynamics between R-domain and C42 in vascular KATP channels

Molecular dynamics between ED-domain and KNt in vascular KATP channels

## Acknowledgements

A portion of this research was supported by NIH grant U24GM129547 and performed at the Pacific Northwest Cryo-EM Center (PNCC) at Oregon Health & Science University and accessed through EMSL (grid.436923.9), a DOE Office of Science User Facility sponsored by the Office of Biological and Environmental Research. We acknowledge support by the National Institutes of Health grants R01GM145784 and R35GM161749 (to SLS), an Oregon Research Medical Foundation Career Development Award (to CMD), an American Heart Association Postdoctoral scholar award 25POST1373218 (to YYK). We acknowledge the funding by the National Institutes of Health grant R35HL171542 (to CGN). We acknowledge Polish high-performance computing infrastructure PLGrid for awarding this project access to the LUMI supercomputer, owned by the EuroHPC Joint Undertaking, hosted by CSC (Finland) and the LUMI consortium through PLL/2025/09/018870 (to KWS).

## Author contributions

CMD designed and performed cryoEM experiments, prepared samples, analyzed data, prepared figures, wrote and edited the manuscript. YYK prepared cryoEM samples, performed electrophysiology experiments, analyzed data, prepared figures, and edited the manuscript. ZY prepared samples. SJL prepared samples and performed cryoEM experiments. KWS performed MD simulation experiments, analyzed data, prepared figures, wrote and edited the manuscript. SLS conceived the project, analyzed data, prepared figures, wrote and edited the manuscript. CGN conceived the project and edited the manuscript.

## Abbreviations

K_ATP_: ATP-sensitive potassium channel
Kir: inward-rectifying potassium channel
SUR1: sulfonylurea receptor 1
CTD: cytoplasmic domain of Kir6
KNt: Kir6 N-terminal tail
TMD: transmembrane domain
NBD: nucleotide binding domain of SUR
N1-T2: nucleotide binding domain 1-transmembrane domain 2 of SUR
ECL: extracellular loop
ICL: intracellular loop
IDR: intrinsically disordered region
M1 or M2: membrane helix 1 or 2 of Kir6.1
cryoEM: cryogenic electron microscopy
TM: transmembrane
ATP: adenosine triphosphate
ADP: adenosine diphosphate
PIP_2_: phosphatidylinositol-4,5-bisphosphate

