## Supplemental Figures for "MgATP/MgADP-dependent conformational dynamics and intrinsically disordered regions of vascular K_ATP_ channels revealed by cryoEM"

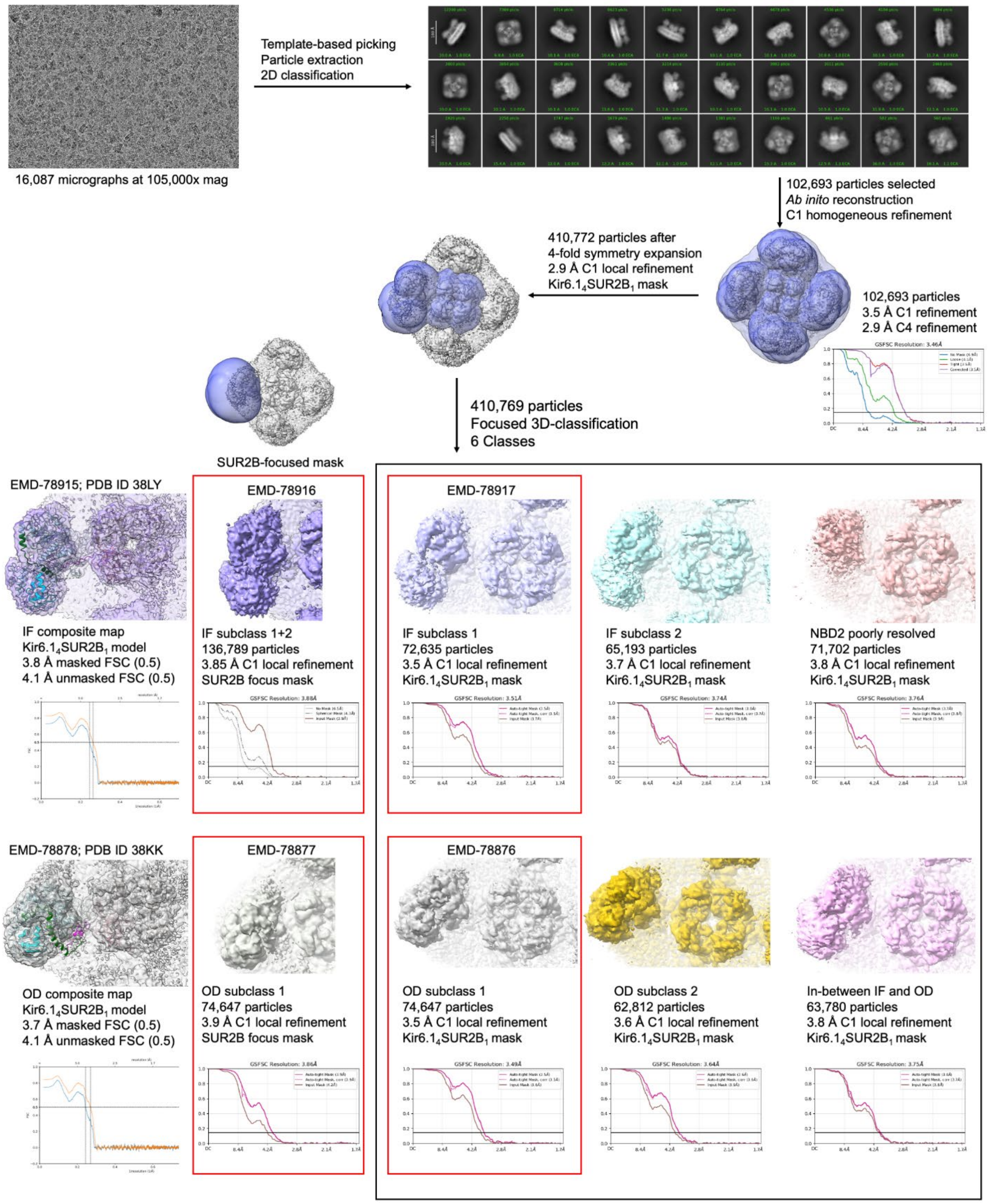
16,087 micrographs at 105,000x mag
Template-based picking
Particle extraction
2D classification
102,693 particles selected
*Ab inito* reconstruction
C1 homogeneous refinement
410,772 particles after
4-fold symmetry expansion
2.9 Å C1 local refinement
Kir6.$1_4$SUR2$B_1$ mask
102,693 particles
3.5 Å C1 refinement
2.9 Å C4 refinement
GSFSC Resolution: 3.46Å
410,769 particles
Focused 3D-classification
6 Classes
SUR2B-focused mask
EMD-78915; PDB ID 38LY
IF composite map
Kir6.$1_4$SUR2$B_1$ model
3.8 Å masked FSC (0.5)
4.1 Å unmasked FSC (0.5)
EMD-78916
IF subclass 1+2
136,789 particles
3.85 Å C1 local refinement
SUR2B focus mask
GSFSC Resolution: 3.88Å
EMD-78917
IF subclass 1
72,635 particles
3.5 Å C1 local refinement
Kir6.$1_4$SUR2$B_1$ mask
GSFSC Resolution: 3.51Å
IF subclass 2
65,193 particles
3.7 Å C1 local refinement
Kir6.$1_4$SUR2$B_1$ mask
GSFSC Resolution: 3.74Å
NBD2 poorly resolved
71,702 particles
3.8 Å C1 local refinement
Kir6.$1_4$SUR2$B_1$ mask
GSFSC Resolution: 3.76Å
EMD-78878; PDB ID 38KK
OD composite map
Kir6.$1_4$SUR2$B_1$ model
3.7 Å masked FSC (0.5)
4.1 Å unmasked FSC (0.5)
EMD-78877
OD subclass 1
74,647 particles
3.9 Å C1 local refinement
SUR2B focus mask
GSFSC Resolution: 3.86Å
EMD-78876
OD subclass 1
74,647 particles
3.5 Å C1 local refinement
Kir6.$1_4$SUR2$B_1$ mask
GSFSC Resolution: 3.49Å
OD subclass 2
62,812 particles
3.6 Å C1 local refinement
Kir6.$1_4$SUR2$B_1$ mask
GSFSC Resolution: 3.64Å
In-between IF and OD
63,780 particles
3.8 Å C1 local refinement
Kir6.$1_4$SUR2$B_1$ mask
GSFSC Resolution: 3.75Å

**Figure S1.** Data processing workflow to classify and resolve distinct SUR2B-NBD-dimerized (EMD-78878; PDB ID 38KK) and SUR2B-NBD-separated (EMD-78915; PDB ID 38LY) structures of the $K_{ATP}$ channel from samples prepared in the presence of MgADP/MgATP. After patch motion-correction and patch CTF-estimation in cryoSPARC, 16,087 micrographs were used to pick particles. After extraction, $K_{ATP}$ channel particles were classified into 2D classes that showed obvious conformational heterogeneity of the SUR2B subunits. Particles selected from good 2D classes and without symmetry imposed were used to reconstruct a 3.5 Å cryoEM map. These particles were 4-fold symmetry expanded and subsequent refinements used a focus map covering the Kir6.1 core plus one SUR2B subunit. Particles were classified into distinct populations using a focused 3D classification strategy. Six main 3D classes were resolved, two classes in an NBD-separated inward-facing conformation with the NBDs further out away from Kir6.1-CTD or closer in towards Kir6.1-CTD (IF subclass 1 or IF subclass 2, blue and teal maps, respectively), two classes in an SUR2B- NBD-dimerized occluded (OD) conformation with the NBDs closer in towards Kir6.1-CTD or further out away from Kir6.1-CTD (OD subclass 1 or OD subclass 2, grey or yellow maps, respectively), a poorly resolved NBD2 class (peach map), and a dynamic NBD2 class (pink map). Further local refinement of the IF-out class and the OD-in class was carried out using a SUR2B without TMD0 mask. The Kir6.1 tetramer was in a CTD-down (inactivated) closed-conformation bound to ATP in all classes. All maps are displayed at a 2.5 rmsd contour.

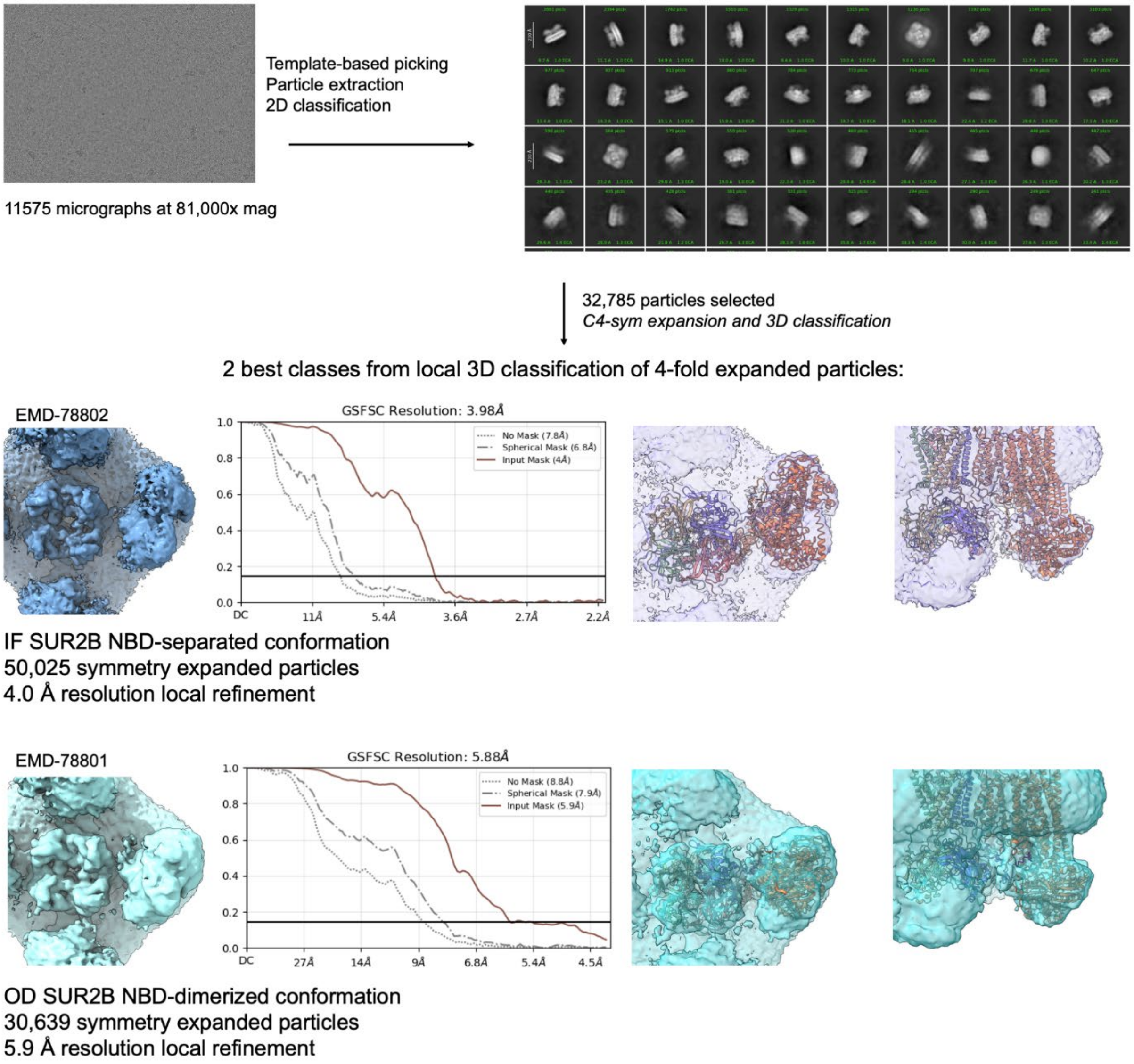


**Figure S2.** Data processing workflow to classify SUR2B-NBD-dimerized (OD) and SUR2B-NBD-separated (IF) conformations of the $vK_{ATP}$ channel from samples prepared with MgADP/MgATP but without VU0542270. After patch motion-correction and patch CTF-estimation in cryoSPARC, 11,575 micrographs were used to pick particles. After extraction, $K_{ATP}$ channel particles were classified into 2D classes. Particles were selected from good 2D classes and were 4-fold symmetry expanded and subsequent refinements used a focus map covering the Kir6.1 core plus one SUR2B subunit. Particles were classified into an IF conformation (blue map) and an OD conformation (cyan map) using a focused 3D classification strategy. The structural models of the IF and OD conformations (PDB ID 38LY and PDB ID 38KK, respectively; see Figure S1) are fit as a ridged body into each map.

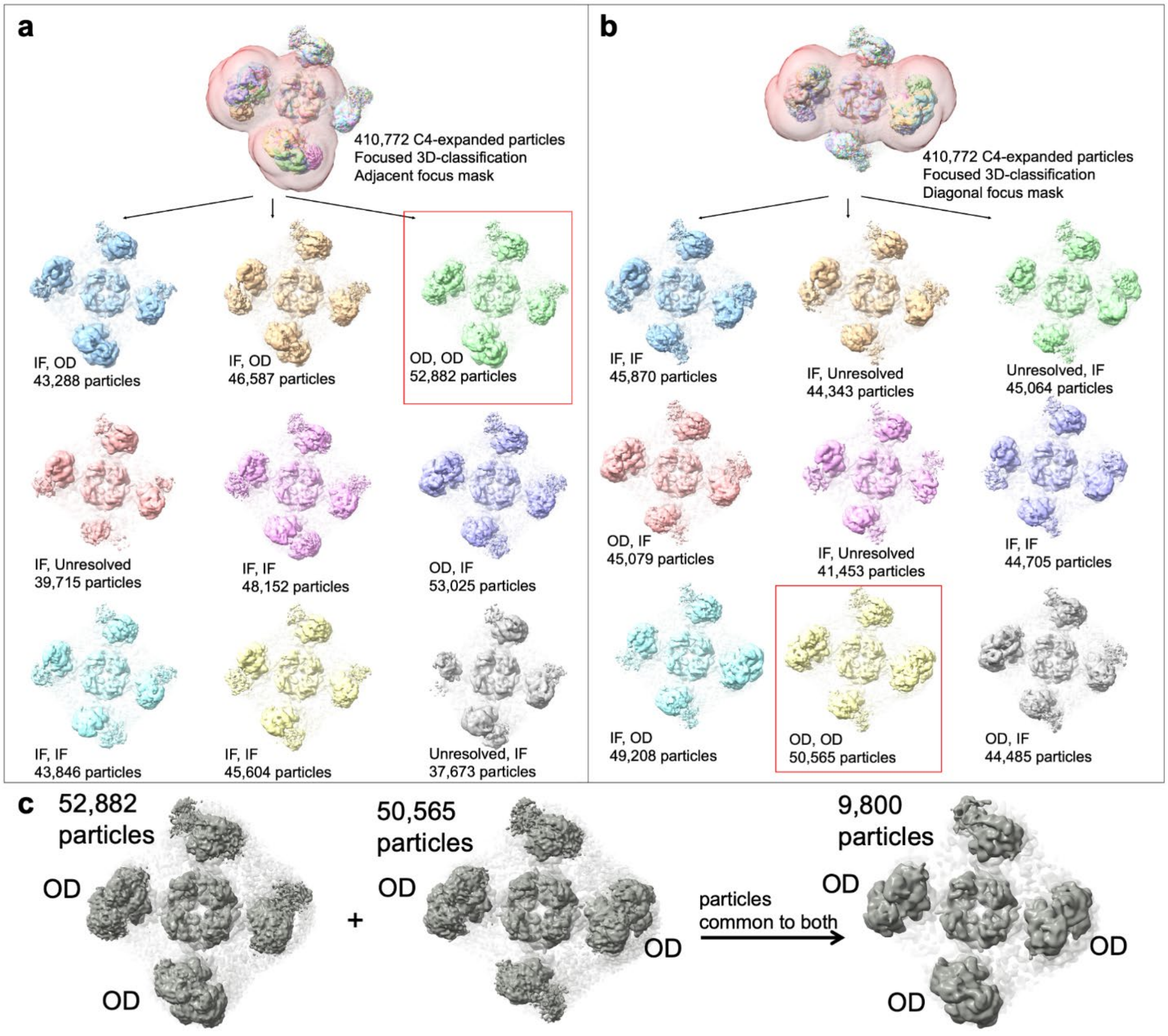


**Figure S3.** Vascular $K_{ATP}$ channels with multiple SUR2B subunits in an occluded NBD-dimerized (OD) conformation. **(a)** 3D classification of 410,772 symmetry expanded particles into 9 classes with the focus map of the Kir6.1 tetramer and adjacent SUR2B subunits shown in semi-transparent red. The cryoEM map of both adjacent SUR2B subunits in an NBD-dimerized OD conformation is shown in green inside a red box, with the conformation of both SUR2B subunits in other classes listed below each map along with the number of particles. **(b)** 3D classification focused on the Kir6.1 tetramer and diagonal SUR2B subunits (semi-transparent red focus mask). The cryoEM map of both diagonal SUR2B subunits in an NBD-dimerized OD conformation is shown in yellow inside a red box. **(c)** The particle numbers and maps of both the adjacent and diagonal SUR2Bs in the OD conformation are shown as grey maps with the NBD-dimerized SUR2B subunits labeled (OD) and the poorly resolved SUR2B subunits not labeled. A local refinement of the 9,800 particles common to both has three SUR2B subunits clearly in the OD conformation, and would include particles that have all four SUR2B subunits in the OD conformation, but low particle number prevented accurate further 3D classification.

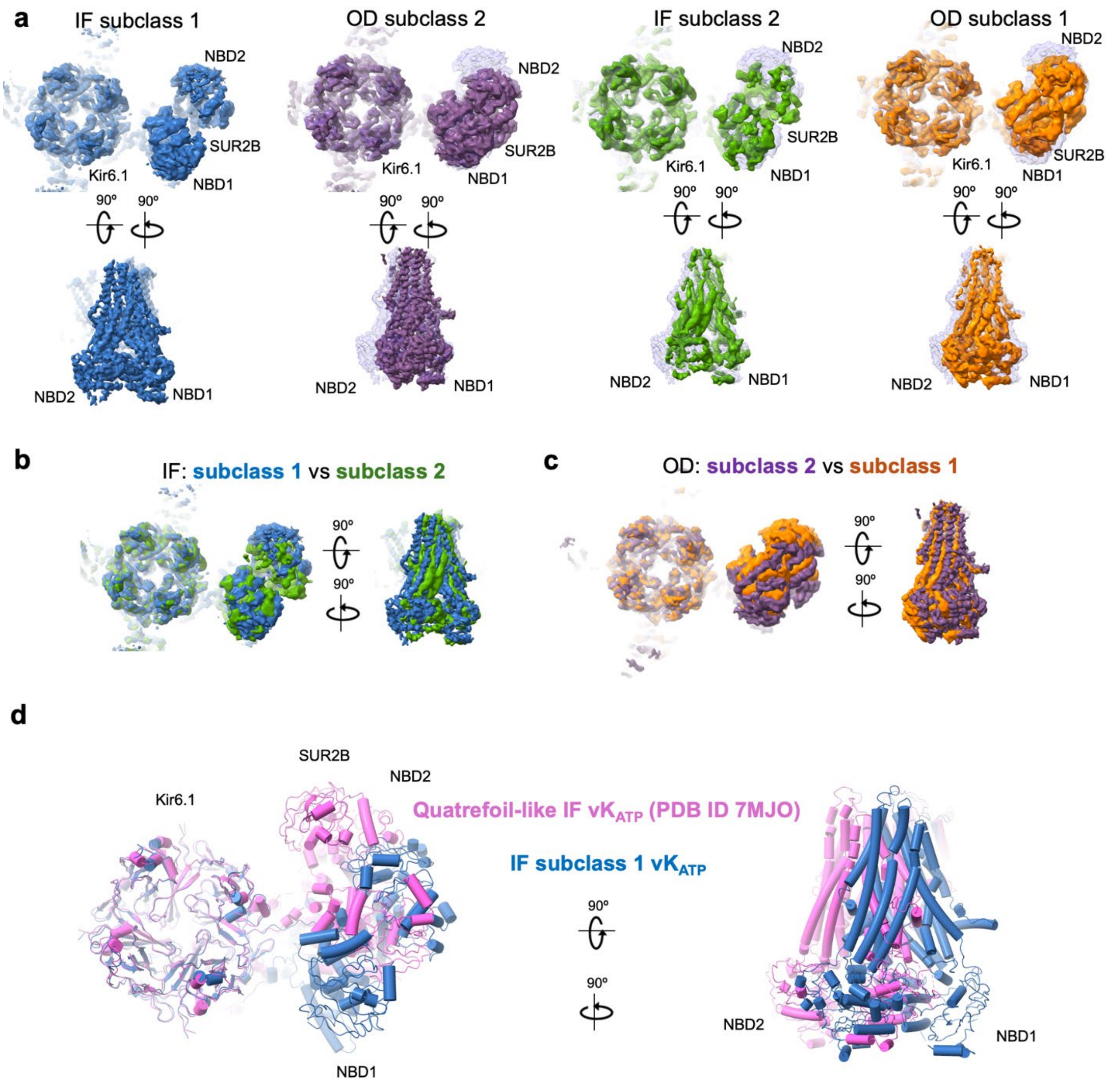


**Figure S4. Structural dynamics of the SUR2B subunit in $vK_{ATP}$ channels. (a)** Comparison between the subclasses of $vK_{ATP}$ channels in NBD-dimerized OD conformations and NBD-separated IF conformations. CryoEM maps for the IF conformation subclass 1 (blue) and the OD conformation subclass 2 (purple) are shown first on the left as these are the two main subclasses used to build models for structural analysis (see Results in the main text). These are followed by the IF conformation subclass 2 (green) and OD conformation subclass 1 (orange). All maps are overlaid on a grey transparent map of the IF conformation subclass 1. Views are from the cytoplasm and the side. **(b)** Overlay of cryoEM maps of the two IF conformation subclasses in (a) showing that in subclass 2, the SUR2B-ABC core is closer to the Kir6.1-CTD. **(c)** Overlay of cryoEM maps of the two OD conformation subclasses, with subclass 2 showing the SUR2B-ABC core closer to the Kir6.1-CTD. **(d)** Comparison of the IF conformation subclass 1 in this study with our previously published quatrefoil-like $vK_{ATP}$ channel structure bound to inhibitory ATP and glibenclamide (pink; PDB ID 7MJO). To emphasize the overall conformational differences of SUR2B relative to Kir6.1 in the structures, all maps are shown at a high contour of 0.10 V (~6 rmsd) at which only the most ordered parts of the structures are visible with noise from the micelle and more disordered regions hidden.

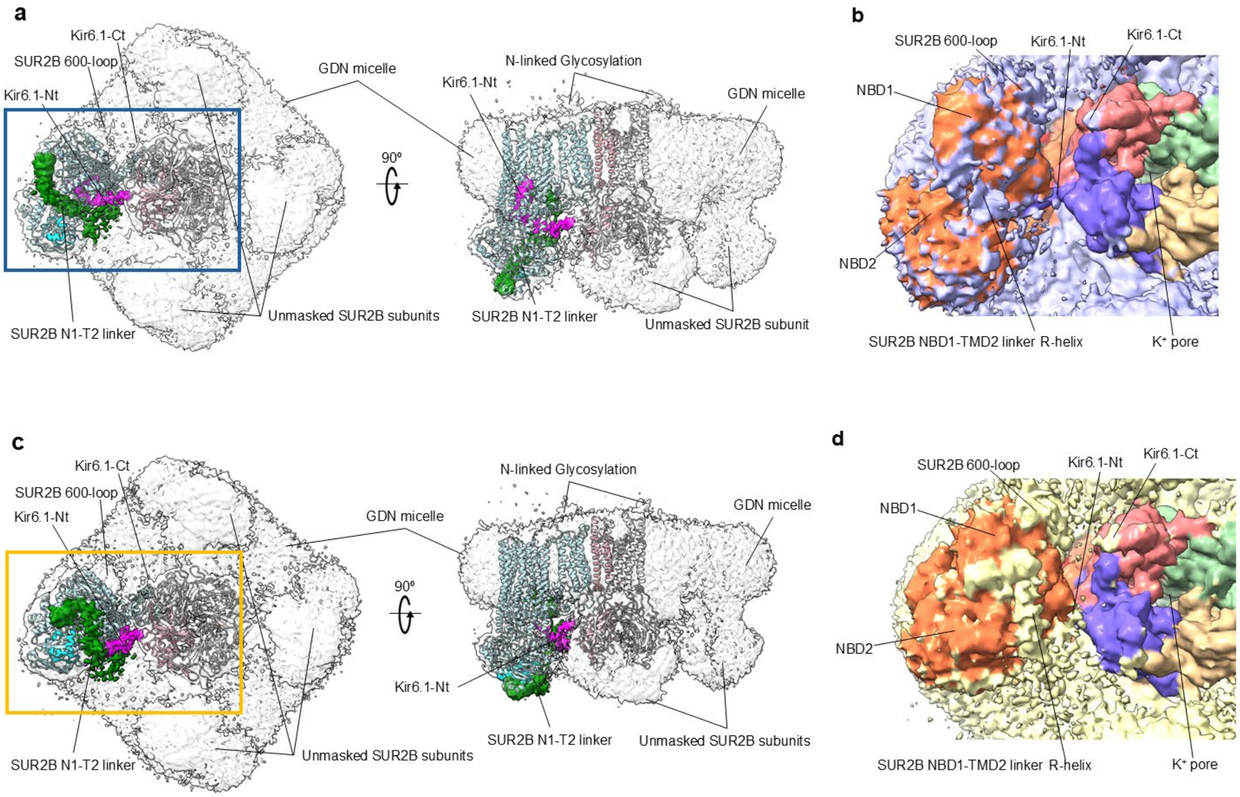


**Figure S5.** Full CryoEM density reconstruction from symmetry-expanded local refinements with a mask to focus on one SUR2B subunit plus the Kir6.1 tetramer core in both the NBD-dimerized OD and NBD-separated IF major conformations. The unmasked map regions include three SUR2B subunits and the Glyco-diosgenin micelle. Intrinsically disordered regions of the $K_{ATP}$ channel include the Kir6.1 N-terminal (Kir6.1-Nt) and C-terminal (Kir6.1-Ct), and the SUR2B 600-loop (corresponding to TMD1-NBD1 linker) and NBD1-TMD2 linker (N1-T2 linker). **(a)** CryoEM map of the NBD-separated IF conformation is shown at 2.5 rmsd (0.045 V), colored zones are shown within 5.0 Å of the defined regions of the model. Kir6.1 residues 1-32 make up the Kir6.1-Nt and are colored magenta, and remaining residues of one Kir6.1 subunit are colored light pink. SUR2B residues 908-974 make up the N1-T2 linker and are colored green. The SUR2B C-terminal 42 residues (C42; residues 1503-1545) that are unique to SUR2B are colored cyan. **(b)** Cytoplasmic view focused on the box region in panel (a) of the map of the $vK_{ATP}$ channel in the IF conformation that is colored uniquely for each chain of the IF model but without a structural model for the N1-T2 linker, the KNt, or other IDRs, with map regions that are not interpreted by the model colored blue. **(c)** The NBD-dimerized OD conformation map is shown at 2.5 rmsd (0.048 V), colored zones are shown within 5.0 Å of the defined regions of the model, colored the same as in (a). **(d)** CryoEM map of the $vK_{ATP}$ channel in the NBD-separated IF conformation, colored by chain but with IDRs removed, with uninterpreted regions colored yellow.

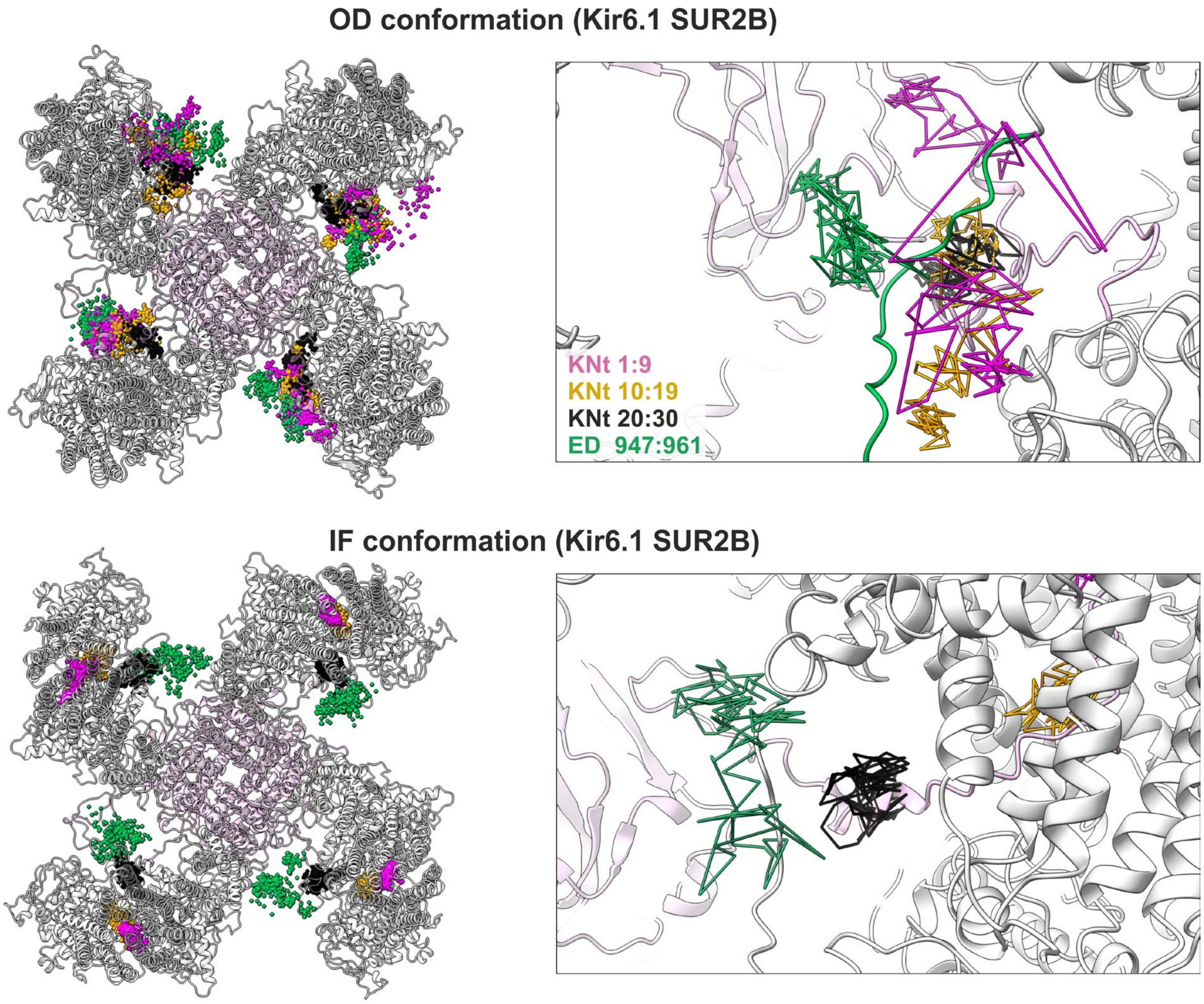


**Figure S6.** Time evolution of the centers of mass of selected intrinsically disordered regions during MD simulations. The left panels show trajectories from all MD simulations. The right panels show two representative trajectories for each IDR (same color as in the left panels) to illustrate the spatial evolution of individual simulations more clearly.

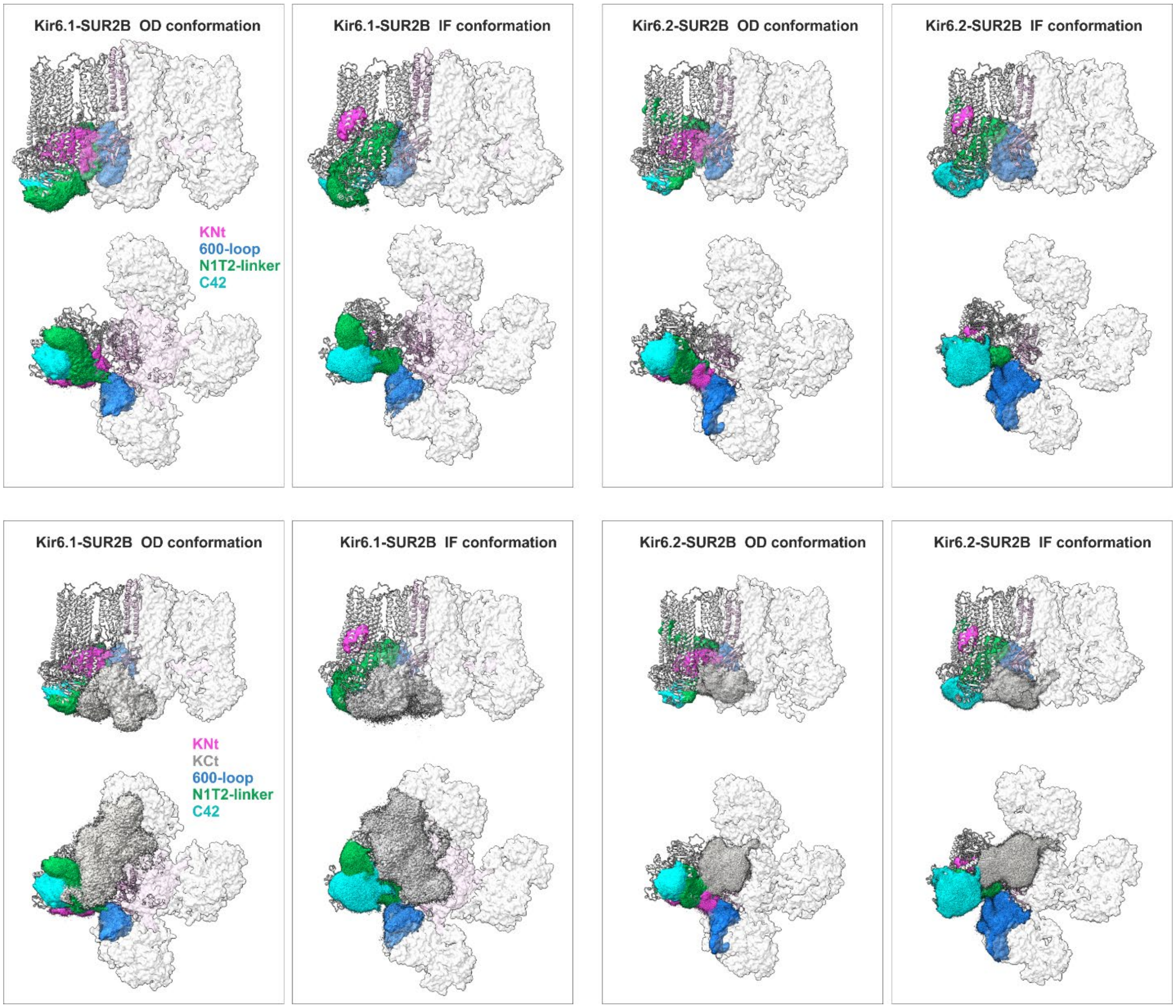


**Figure S7.** Volumes sampled by selected $vK_{ATP}$ IDRs during MD simulations. The displayed volumes represent the spatial regions explored by each IDR over the course of the simulations.

### Table S1. Map and model statistics of SUR2B/Kir6.1 $K_{ATP}$ channels

| | NBD-separated inward-facing $Kir6.1_4SUR2B_1$ EMD-78917 | NBD-separated inward-facing SUR2B focus EMD-78916 | NBD-dimerized occluded $Kir6.1_4SUR2B_1$ EMD-78876 | NBD-dimerized occluded SUR2B focus EMD-78877 | IF $Kir6.1_4SUR2B_1$ (no VU270) EMD-78802 | OD $Kir6.1_4SUR2B_1$ (no VU270) EMD-78801 |
|---|---|---|---|---|---|---|
| **Data Collection** | | | | | | |
| Microscope | Titan Krios | | | | Titan Krios | |
| Voltage (kV) | 300 | | | | 300 | |
| Camera | Gatan K3 Biocontinuum | | | | Gatan K3 Biocontinuum | |
| Camera mode | Super-resolution | | | | Super-resolution | |
| Defocus range (µm) | -0.5 ~ -2.5 | | | | -0.5 ~ -2.5 | |
| Movies | 16,087 | | | | 11,575 | |
| Frames/movie | 70 | | | | 70 | |
| Exposure time (s) | 3.9 | | | | 3.9 | |
| Frame rate (/s) | 17.9 | | | | 17.9 | |
| Magnified pixel size (Å)* | 0.827 (0.4135 super-res) | | | | 1.059 (0.5295 super-res) | |
| Total dose ($e^-/Å^2$) | 55 | | | | 55 | |
| **Reconstruction** | | | | | | |
| Software | CryoSPARC v5 BETA | CryoSPARC v5 BETA | CryoSPARC v5 BETA | CryoSPARC v5 BETA | CryoSPARC v5 BETA | CryoSPARC v5 BETA |
| Symmetry | C1 local refine | C1 local refine | C1 local refine | C1 local refine | C1 local refine | C1 local refine |
| Mask | $Kir6.1_4SUR2B_1$ | $SUR2B_1$ | $Kir6.1_4SUR2B_1$ | $SUR2B_1$ | $Kir6.1_4SUR2B_1$ | $Kir6.1_4SUR2B_1$ |
| Particles (C4-expanded) | 72,635 | 136,789 | 74,647 | 74,647 | 50,025 | 30,639 |
| Resolution (GSFSC=0.143) | 3.5 Å | 3.85 Å | 3.5 Å | 3.9 Å | 4.0 Å | 5.9 Å |
| **Refinement Statistics** | | | | | | |
| PDB ID | 38LY | | 38KK | | | |
| Composite map | EMD-78915 | | EMD-78878 | | | |
| Atoms | 21,484 | | 22,728 | | | |
| Protein residues | 2,848 | | 2,846 | | | |
| Map CC (masked) | 0.82 | | 0.86 | | | |
| FSC model (0.5) | 3.8 Å | | 3.7 Å | | | |
| Clash score | 5.43 | | 6.65 | | | |
| Molprobity score | 1.89 | | 2.24 | | | |
| Cβ deviations | 0 | | 0 | | | |
| Rotamer outliers (%) | 1.99 | | 6.11 | | | |
| ADP (mean protein) | 126.17 | | 136.75 | | | |
| ADP (mean ligands) | 126.27 | | 133.77 | | | |
| **Ramachandran** | | | | | | |
| Outliers | 0 | | 0 | | | |
| Allowed | 5.40 | | 3.99 | | | |
| Favored | 94.60 | | 96.01 | | | |
| **Bonds (RMSD)** | | | | | | |
| Length (Å) | 0.002 | | 0.003 | | | |
| Bond angles | 0.444 | | 0.475 | | | |
